# Lipids Regulate Export of Lysosomal Enzymes from the Endoplasmic Reticulum

**DOI:** 10.64898/2026.04.16.719038

**Authors:** Baolong Xia, Myeonghoon Han, Isaac Park, Norbert Perrimon

## Abstract

Lysosomal enzymes are synthesized in the Endoplasmic Reticulum (ER) and transported to lysosomes to execute their functions. Deficiencies in lysosomal enzymes or components of the lysosomal transport machinery result in lysosomal storage disorders. While mannose-6-phosphate mediated lysosomal enzymes sorting in the Golgi has been extensively characterized, the mechanisms governing their export from the ER remain elusive. Here, we show that de novo lipogenesis, a metabolic pathway responsible for fatty acid synthesis, regulates lysosomal enzyme transport. Inhibition of de novo lipogenesis leads to the retention of lysosomal enzymes within the ER. Mechanistically, fatty acid derived from de novo lipogenesis is used for Arf1 myristoylation. Myristoylated Arf1 promotes retrograde vesicle trafficking from the Golgi to the ER, thereby maintaining the homeostatic bidirectional flux required for efficient ER export of lysosomal enzymes. Our findings uncover a critical functional link between lipid metabolism and lysosomal enzyme trafficking.

**Significance:** Proper intracellular protein trafficking is essential for cellular homeostasis, yet how cellular metabolism regulates this process remains poorly understood. In this study, we show that inhibition of fatty acid synthesis disrupts lysosomal enzyme trafficking by causing these enzymes to accumulate in the endoplasmic reticulum instead of reaching lysosomes. Fatty acids are required for protein lipidation that supports vesicle trafficking and export of lysosomal enzymes from the endoplasmic reticulum. In addition, interactome screening identified other regulators of lysosomal enzyme transport. These findings reveal a direct link between fatty acid metabolism and lysosomal protein trafficking, expanding our understanding of how metabolic pathways coordinate intracellular transport processes.

## Introduction

Lysosomes are membrane-bound organelles responsible for the degradation, processing, and recycling of macromolecules. Their functions are mediated by more than 50 soluble hydrolytic enzymes, including proteases (cathepsins), lipases, glycosidases, nucleases, phosphatases, and sulfatases. These enzymes are synthesized in the endoplasmic reticulum (ER) and subsequently transported to lysosomes. Deficiencies in lysosomal enzymes or components of the lysosomal transport machinery lead to lysosomal storage disorders, which are characterized by the accumulation of undegraded substrates due to impaired lysosomal function.

Mannose-6-phosphate (M6P) dependent post-Golgi trafficking is a well-characterized pathway that mediates the delivery of lysosomal enzymes from the Golgi apparatus to lysosomes in mammalian cells. In the Golgi, lysosomal enzymes are modified with M6P and recognized by M6P receptors for transport to lysosomes. The generation of M6P is catalyzed sequentially by two enzyme complexes: N-acetylglucosamine (GlcNAc)-1-phosphotransferase (GNPT) and GlcNAc-1-phosphodiester α-N-acetylglucosaminidase (also known as the uncovering enzyme)(1). GNPT recognizes lysosomal enzymes and transfers the GlcNAc-1-phosphate moiety from UDP-GlcNAc to mannose residues on the enzymes. The uncovering enzyme subsequently removes the terminal GlcNAc, exposing a phosphate group linked to mannose, thereby enabling binding to M6P receptors. GNPT is a hexameric complex composed of α_2_β_2_γ_2_subunits and is encoded by two genes: *GNPTAB*, which encodes the α/β precursor, and *GNPTG*, which encodes the γ subunit. Proteolytic activation of the α/β precursor is mediated by site-1 protease (S1P)(2). Notably, *Drosophila* lacks discernible homologs of GNPTG and uncovering enzyme required for M6P modification. Moreover, the *Drosophila* M6P receptor ortholog Lerp lacks amino acids implicated in M6P binding, and consistent with this, Lerp does not bind to phosphomannan affinity column(3, 4). Whether and how S1P is involved in lysosomal enzyme trafficking in this scenario is unclear.

In addition to its role in GNPTAB activation, S1P plays a well-established role in lipid metabolism through proteolytic cleavage of sterol regulatory element-binding proteins (SREBPs), transcription factors that regulate lipid synthesis and cholesterol homeostasis(5). In mammalian cells, under conditions of low cholesterol, SREBP is translocated from the ER to the Golgi apparatus, where it undergoes sequential cleavage by S1P and site-2 protease (S2P). This process releases the N-terminal transcriptional activation domain to the nucleus to activate target genes involved in lipid metabolism. In *Drosophila*, however, SREBP translocation from the ER to the Golgi is regulated by fatty acid palmitate rather than sterols(6). Moreover, S2P cleavage is not absolutely required for SREBP activation in flies, as alternative proteases, such as the caspase Drice, can process SREBP in the absence of S2P(7).

Although the role of S1P in lysosome biogenesis and function through proteolytic activation of GNPTAB has been well established, it remains unclear whether its function in lipid metabolism also contributes to lysosomal regulation. Several indirect lines of evidence have complicated this interpretation(2, 8, 9). For instance, in zebrafish, knockdown of either *S1P* or *S2P* results in both lipid abnormalities and cartilage defects, whereas knockdown of *SREBP cleavage-activating protein* (*SCAP*), which is essential for SREBP processing, results only in strong lipid phenotypes with normal cartilage(8).Based on these observations, it has been proposed that the cartilage defects observed upon *S1P* depletion occur independently of the lipid abnormalities caused by impaired SCAP-mediated SREBP activation. However, it is not clear how *S2P* knockdown produces similar cartilage phenotypes, leaving the underlying mechanism unresolved.

In this study, we provide direct evidence that the *S1P-SREBP* axis plays a critical role in lysosomal enzyme trafficking in *Drosophila* S2R+ cells. Knockdown of *SREBP* results in the accumulation of lysosomal enzymes within the ER, indicating a defect in ER export. Mechanistically, fatty acid derived from de novo lipogenesis is used for Arf1 myristoylation. Myristoylated Arf1 facilitates retrograde vesicle trafficking from the Golgi to the ER, thereby maintaining the homeostatic bidirectional flux required for efficient ER export of lysosomal enzymes.

## Results

### SREBP regulates CTSL exiting endoplasmic reticulum (ER)

To investigate the mechanisms underlying lysosomal enzyme transport to the lysosome, we analyzed previously published genome-wide CRISPR screens for lysosomal function(10, 11) and identified *S1P* as one of the top hits. S1P is known to cleave the precursor of GlcNAc-1-phosphotransferase (GNPT), generating mature GNPT subunits required for mannose-6-phosphate modification of lysosomal enzymes(2). However, whether SREBP, another substrate of S1P(5) (Figure 1A), participates in lysosomal enzyme trafficking remains unclear.

**Figure 1.**
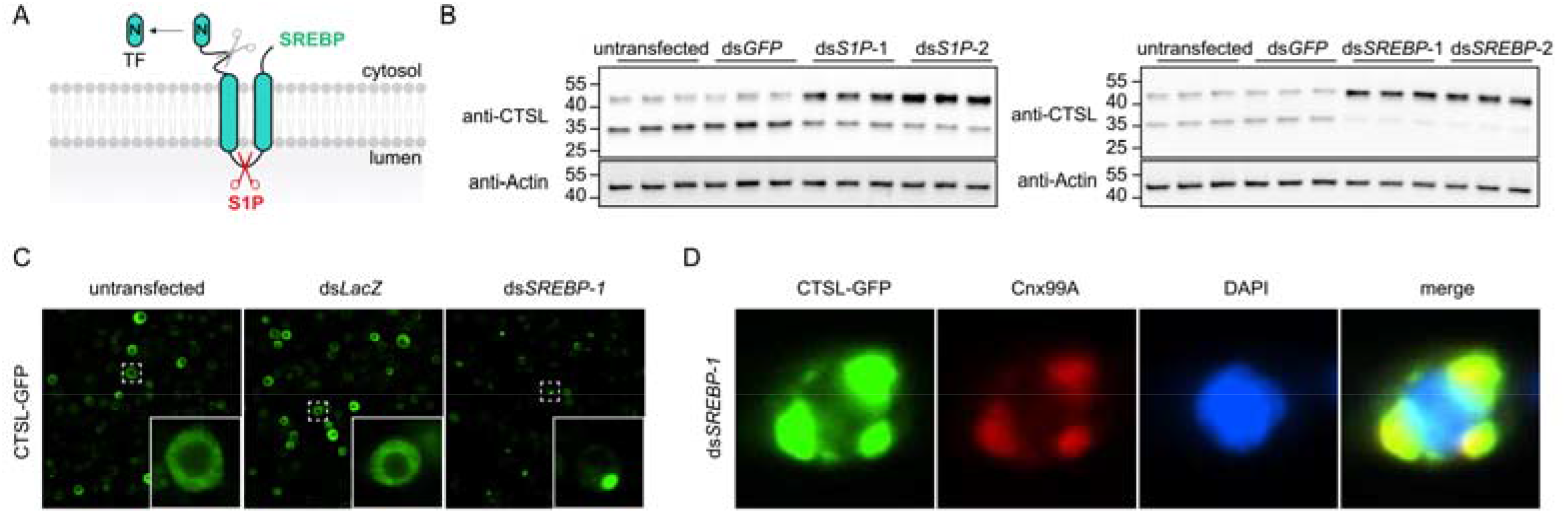
The transcription factor SREBP regulates CTSL export from the ER. A. Schematic of the processing of SREBP by S1P on the Golgi membrane, resulting in release of the N-terminal transcriptional activation domain. B. Western blot analysis of CTSL in the *S1P* and *SREBP* knockdown cells. Two different non-overlapping dsRNAs were used for each gene. The upper band corresponds to the precursor form of CTSL, and the lower band represents the mature form. C. Subcellular localization of CTSL-GFP in cells transfected with dsRNAs targeting *lacZ* (control) or *SREBP*. D. Immunostaining of the ER marker Cnx99A in CTSL-GFP expressing cells following *SREBP* knockdown.

To determine whether the *S1P-SREBP* axis regulates lysosomal enzyme transport, we used two non-overlapping double-stranded RNAs (dsRNAs) to deplete *S1P* or *SREBP* in *Drosophila* S2R+ cells and examined the lysosomal protease cathepsin L (CTSL). CTSL is synthesized in the endoplasmic reticulum as a precursor and subsequently processed into its mature form upon delivery to the lysosome. Thus, processing of the precursor form serves as a functional readout of CTSL trafficking(3). Notably, compared with untransfected cells or cells transfected with ds*GFP*, depletion of either *S1P* or *SREBP* led to accumulation of the precursor form of CTSL (Figure 1B), indicating impaired CTSL transport.

To determine the effects of *SREBP* depletion on CTSL localization, we introduced *GFP* gene into the endogenous *CTSL* locus to express CTSL-GFP fusion protein with GFP fused to the C-terminus of CTSL. This strategy enabled visualization of CTSL subcellular distribution by monitoring GFP fluorescence in cells. In untransfected cells or cells transfected with ds*LacZ*, the GFP signal was diffusely distributed throughout the endomembrane system within the cytoplasm. In contrast, *SREBP* knockdown resulted in a marked redistribution of GFP, characterized by the formation of prominent intracellular puncta (Figure 1C). Moreover, the GFP puncta were colocalized with the ER marker Cnx99A (Figure 1D), indicating that CTSL is retained in the ER in *SREBP*-depleted cells. This observation is consistent with the accumulation of the precursor form of CTSL detected by immunoblotting upon *SREBP* knockdown.

### Fatty acids regulate CTSL transport through protein myristoylation

Next, we investigated the mechanism by which the transcription factor *SREBP* regulates lysosomal enzyme transport. Given that *SREBP* functions as a master regulator of de novo lipogenesis(12), we hypothesized that this metabolic pathway contributes to lysosomal enzyme trafficking. De novo lipogenesis is a metabolic process through which fatty acids are synthesized from carbohydrates. The pathway is initiated by the generation of acetyl-CoA from citrate via ATP citrate lyase (ACLY) and from acetate via acetyl-CoA synthase (ACS). Acetyl-CoA is subsequently carboxylated by acetyl-CoA carboxylase (ACC) to produce malonyl-CoA. In the final step, fatty acid synthase (FASN1) catalyzes successive condensation reactions, utilizing malonyl-CoA as a two-carbon donor and acetyl-CoA as a primer, ultimately generating long-chain fatty acids (Figure 2A).

**Figure 2.**
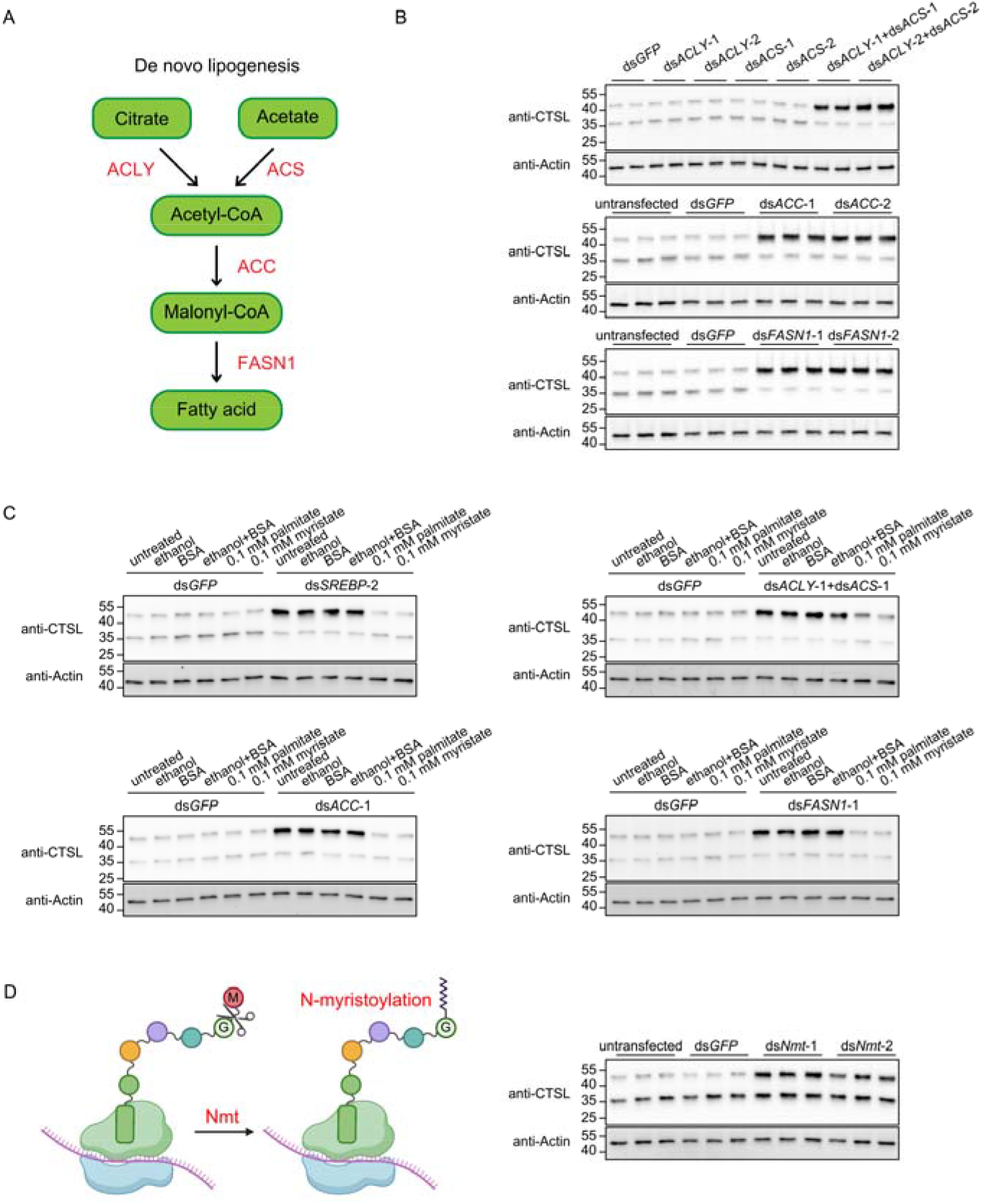
De novo lipogenesis regulates CTSL transport through protein myristoylation. A. Schematic of the de novo lipogenesis pathway. Key enzymes involved in each step are highlighted in red: ACLY (ATP citrate lyase), ACS (acetyl-CoA synthase), ACC (acetyl-CoA carboxylase), FASN1 (fatty acid synthase). B. Western blot analysis of CTSL following knockdown of individual enzymes in the lipogenesis pathway. Two different non-overlapping dsRNAs targeting each gene were used. C. Western blot analysis of CTSL in dsRNA transfected cells supplemented with 0.1 mM palmitate (C16:0) or myristate (C14:0). D. Western blot analysis of CTSL in *Nmt* knockdown cells. Two different non-overlapping dsRNAs targeting *Nmt* were used. A schematic illustrates the function of N-myristoyltransferase.

We observed that inhibition of any individual steps within the de novo lipogenesis pathway led to accumulation of the precursor form of CTSL, indicating defective lysosomal enzyme transport (Figure 2B). These findings suggest that fatty acid synthesis is required for proper lysosomal trafficking of CTSL. To determine whether the observed phenotype upon blockade of de novo lipogenesis was attributable to fatty acid insufficiency, we examined whether supplementation with end products of the pathway could rescue the accumulation of CTSL precursor in knockdown cells. Palmitate (C16:0) is the primary end product of de novo lipogenesis, with smaller amounts of myristate (C14:0) and stearate (C18:0) also produced(13). Fatty acids were dissolved in ethanol and conjugated to bovine serum albumin (BSA) prior to cellular supplementation. Notably, palmitate or myristate supplementation, but not control treatments, fully rescued the accumulation of the precursor form of CTSL caused by *SREBP* depletion or inhibition of the de novo lipogenesis pathway (Figure 2C). Collectively, these results demonstrate that fatty acids are required for proper lysosomal enzyme transport.

We next sought to determine how these fatty acid metabolites regulate lysosomal enzyme trafficking. Saturated long-chain fatty acids derived from de novo lipogenesis can be further processed into very-long-chain fatty acids by elongases, converted into unsaturated fatty acids by desaturases, incorporated into triglycerides, or utilized for protein acylation by acyltransferases. To dissect which downstream pathway is involved, we used dsRNAs to deplete elongases (SI Appendix, fig. S1A), desaturases (SI Appendix, fig. S1B), or enzymes required for triglyceride synthesis (SI Appendix, fig. S1C) in *Drosophila* S2R+ cells. None of them resulted in accumulation of precursor form of CTSL, suggesting that fatty acid elongation, desaturation, and triglyceride synthesis are dispensable for CTSL trafficking.

In contrast, depletion of acyltransferases led to the accumulation of CTSL precursor (SI Appendix, fig. S1D). One of the hits was *Nmt*, the sole N-myristoyltransferase in *Drosophila*, whose depletion resulted in a marked accumulation of the precursor form of CTSL (Figure 2D). *Nmt* catalyzes cotranslational protein myristoylation by transferring myristate to an N-terminal glycine residue following removal of the initiator methionine(14). Importantly, supplementation with palmitate or myristate failed to rescue the CTSL precursor accumulation induced by *Nmt* knockdown (SI Appendix, fig. S1E), indicating that *Nmt* functions downstream of fatty acid synthesis. Together, these data demonstrate that protein myristoylation is required for proper lysosomal enzyme transport.

### Arf1 myristoylation-dependent vesicle trafficking regulates CTSL transport

To determine how protein myristoylation regulates CTSL transport, we depleted known CTSL-interacting proteins in *Drosophila* S2R+ cells using dsRNAs. Among the candidates tested, *Arf1* depletion resulted in marked accumulation of the CTSL precursor (Figure 3A and SI Appendix, fig. S2A). Arf1 is a small cytoplasmic GTPase that regulates vesicular trafficking by recruiting cargo-sorting coat proteins to membranes(15). Importantly, Arf1 undergoes N-terminal myristoylation, a modification required for its stable association with Golgi membranes(16–19). At the cis-Golgi, myristoylated Arf1 recruits the COPI coat complex to mediate retrograde transport from the Golgi to the ER, whereas at the trans-Golgi, it promotes the formation of clathrin-coated vesicles (Figure 3B). To define which Arf1-dependent pathway is required for CTSL trafficking, we depleted subunits of the COPI and clathrin complexes. Knockdown of individual COPI subunits, but not clathrin subunits, led to accumulation of the CTSL precursor (Figure 3C), indicating that the COPI complex, rather than the clathrin complex, is essential for CTSL transport.

**Figure 3.**
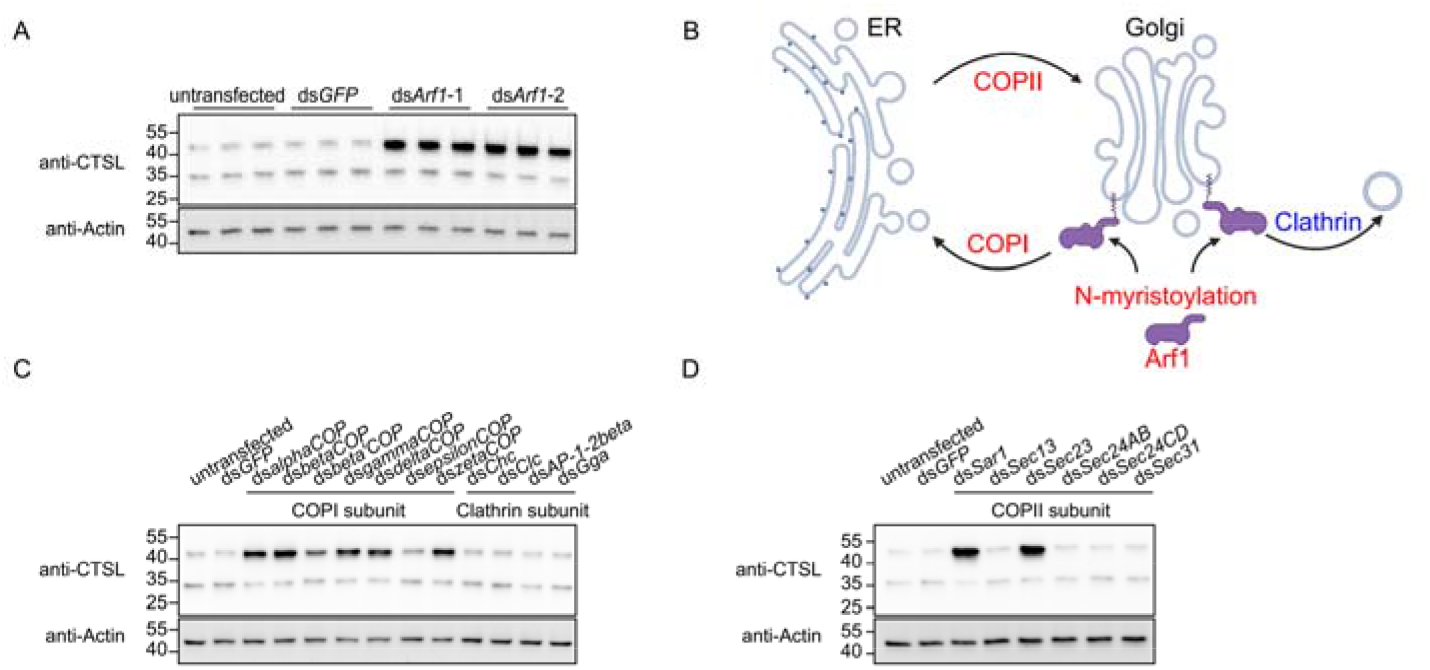
Arf1 myristoylation-dependent vesicle trafficking regulates CTSL transport. A. Western blot analysis of CTSL in *Arf1* knockdown cells. Two different non-overlapping dsRNAs targeting *Arf1* were used. B. Schematic of Arf1 function. Myristoylated Arf1 associates with Golgi membranes to recruit coat proteins for vesicle trafficking. COPI mediates retrograde transport from Golgi to ER, Clathrin mediates post-Golgi transport, COPII mediates anterograde transport from ER to Golgi. C. Western blot analysis of CTSL in cells with knockdown of COPI or clathrin subunits. D. Western blot analysis of CTSL in cells with knockdown of COPII subunits.

Previous genome-wide dsRNA screens in *Drosophila* cells reported markedly increased lipid storage upon depletion of Arf1 or COPI complex components(20, 21), raising the possibility that Arf1/COPI regulates fatty acid homeostasis through lipolysis. To exclude the possibility that CTSL precursor accumulation in Arf1/COPI-depleted cells results from fatty acid insufficiency, we supplemented palmitate or myristate in the knockdown cells. Fatty acid supplementation failed to rescue CTSL precursor accumulation in Arf1- or COPI-depleted cells (SI Appendix, fig. S2B), indicating that the phenotype is not due to impaired fatty acid availability. These results support that Arf1/COPI-mediated retrograde vesicle trafficking, rather than altered lipolysis, is required for CTSL transport.

Vesicular trafficking between the ER and Golgi is coordinated by COPI-mediated retrograde transport and COPII-mediated anterograde transport(22). Given the requirement for COPI-dependent retrograde trafficking in CTSL transport, we next examined whether COPII-mediated anterograde trafficking is also involved. Knockdown of *Sar1* or *Sec23*, the core components of the COPII complex, resulted in accumulation of the CTSL precursor (Figure 3D), suggesting that bidirectional vesicle trafficking between the ER and Golgi is critical for proper CTSL transport.

### Proximity labeling reveals Sccpdh2 as a novel regulator for CTSL transport

To identify additional regulators involved in CTSL transport, we employed a proximity labeling strategy to map novel CTSL-interacting proteins. We fused the engineered biotin ligase TurboID to the C-terminus of CTSL (CTSL-TurboID) to promiscuously biotinylate proximal proteins around CTSL, and used an ER-targeted TurboID protein (BiP-TurboID) that traffics through the endomembrane system as a control (Figure 4A, SI Appendix, fig. S3A). CTSL-TurboID and BiP-TurboID expressing cells exhibited distinct biotinylation patterns (SI Appendix, fig. S3B and S3C), indicating that CTSL possesses a specific proximal interactome distinct from the control. Biotinylated proteins were subsequently isolated using streptavidin beads and identified by mass spectrometry (SI Appendix, Table S1). We then used dsRNAs to deplete proteins enriched in the CTSL-TurboID interactome relative to the BiP-TurboID interactome to assess their roles in CTSL trafficking. Among the candidates tested, depletion of *Tango1* or *Sccpdh2* resulted in accumulation of the CTSL precursor (Figure 4B, SI Appendix, fig. S3D and S3E).

**Figure 4.**
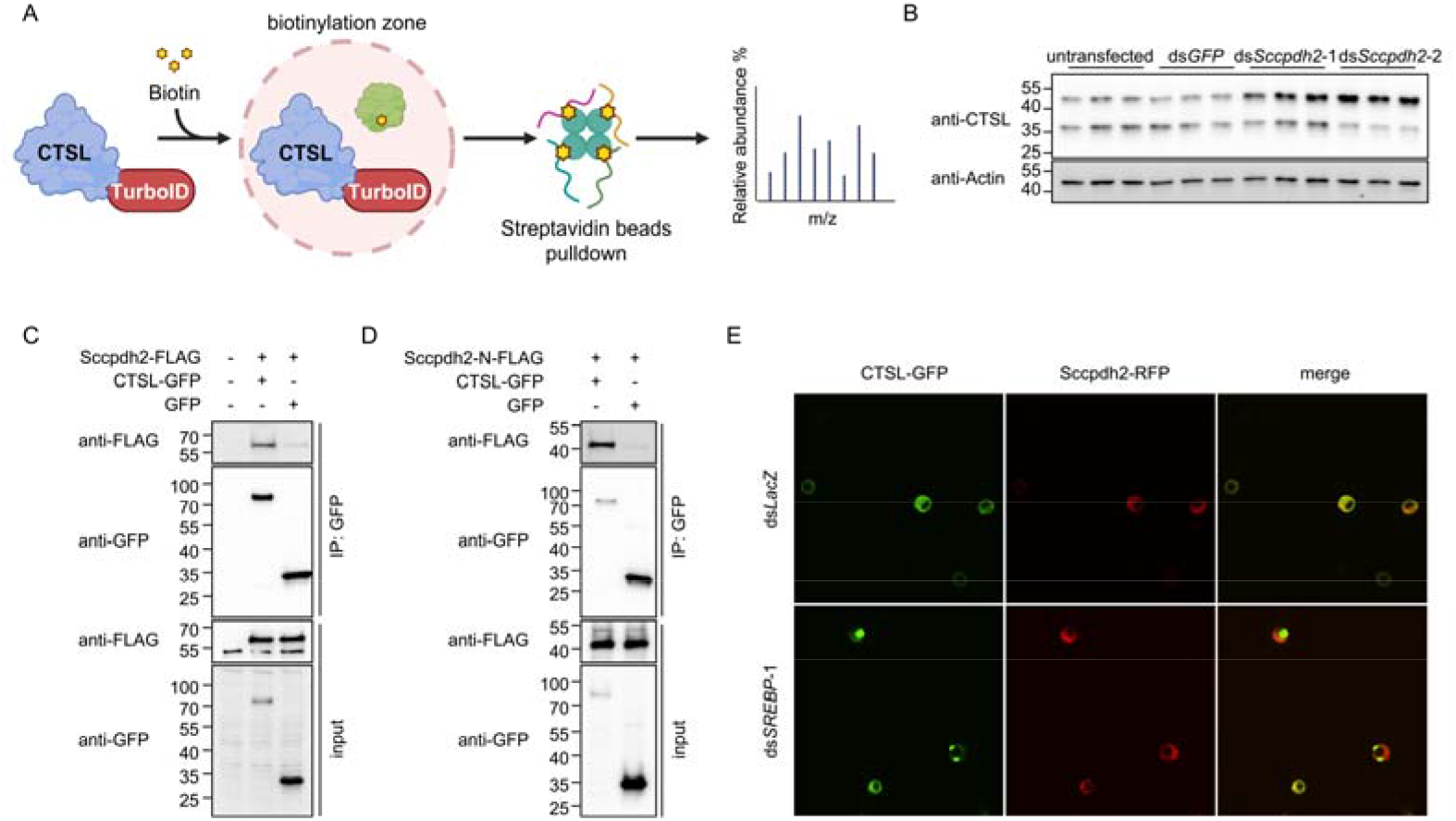
Proximity labelling identifies Sccpdh2 as a novel regulator for CTSL transport. A. Schematic of proximity labeling to biotinylate proteins in proximity to CTSL. B. Western blot analysis of CTSL in *Sccpdh2* knockdown cells. Two different non-overlapping dsRNAs targeting *Sccpdh2* were used. C. Co-immunoprecipitation using GFP-trap beads from lysates of cells co-expressing Sccpdh2-FLAG with CTSL-GFP or GFP control. D. Co-immunoprecipitation using recombinant FLAG-tagged N-terminal fragment of Sccpdh2 incubated with lysates of cells expressing CTSL-GFP or GFP. E. Subcellular localization of CTSL and Sccpdh2 in cells treated with dsRNAs targeting *lacZ* or *SREBP*.

*Tango1* is a well-established regulator of the secretory pathway that organizes membranes and facilitates cargo sorting at ER exit sites(23, 24). In contrast, *Sccpdh2* remains poorly characterized. The human ortholog, *SCCPDH*, exhibits strong co-dependencies in DepMap with the lysosomal trafficking regulator *LYST*, a protein implicated in membrane dynamics and vesicular trafficking(25, 26), suggesting a potential role for *SCCPDH* in lysosomal enzyme transport.

Fatty acid supplementation failed to rescue the accumulation of CTSL precursor in *Sccpdh2* knockdown cells (SI Appendix, fig. S3F), excluding the possibility that Sccpdh2 functions through fatty acids. Since Sccpdh2 was identified through proximity labeling of CTSL, we hypothesized that it physically associates with CTSL. To test this possibility, we performed co-immunoprecipitation in *Drosophila* S2R+ cells using GFP-Trap beads. Compared with the GFP control, Sccpdh2-FLAG was markedly enriched in the CTSL-GFP pulldown sample (Figure 4C), indicating a specific interaction between Sccpdh2 and CTSL.

Topology prediction analysis suggested that *Sccpdh2* encodes a transmembrane protein with a luminal N-terminal domain. To determine whether this luminal domain mediates interaction with CTSL, we incubated a recombinant FLAG-tagged N-terminal fragment of Sccpdh2 with cell lysates expressing either CTSL-GFP or GFP alone. GFP immunoprecipitation assays revealed that the N-terminal domain of Sccpdh2 preferentially associates with CTSL-GFP compared to GFP alone (Figure 4D), indicating that the luminal domain of Sccpdh2 directly interacts with CTSL.

Next, we examined the subcellular localization of Sccpdh2 and CTSL. In ds*LacZ*-treated control cells, Sccpdh2 and CTSL largely colocalized in the endomembrane system within the cytoplasm. However, upon *SREBP* depletion, CTSL formed prominent puncta within the ER, while Sccpdh2 was spatially separated from these puncta (Figure 4E). Taken together, these results support a model in which Sccpdh2 directly interacts with CTSL via its luminal domain and suggest that this interaction is required for proper CTSL trafficking.

## Discussion

In this study, we present that de novo lipogenesis, regulated by the *S1P-SREBP* axis, plays a critical role in lysosomal enzyme trafficking. Fatty acids generated through this pathway are utilized for Arf1 myristoylation. This lipid modification of Arf1 promotes retrograde vesicle trafficking from the Golgi to the ER, thereby sustaining the bidirectional flux necessary for efficient ER export of lysosomal enzymes.

Because ER-to-Golgi transport is essential for the secretion of proteins, we hypothesized that the mechanism identified in this study might more broadly affect ER export of secreted cargo. Consistent with this idea, knockdown of *SREBP*, *Arf1*, or *Sccpdh2* led to intracellular accumulation of the *Drosophila* secreted protein ImpL2 in cell lysates, indicating impaired secretion and consequent retention of ImpL2 within the cell (SI Appendix, fig. S3G). We also observed that Golgi morphology was altered following *SREBP* knockdown. In ds*GFP*-transfected cells, the Golgi marker Golgin84 immunostaining revealed distinct cytoplasmic puncta, consistent with previous reports that the *Drosophila* Golgi apparatus is organized into several discrete, individual stacks. By contrast, in most ds*SREBP*-transfected cells, Golgin84 staining appeared diffuse throughout the cytoplasm, potentially reflecting fusion of Golgi membranes with the ER (SI Appendix, fig. S3H). This phenotype is consistent with prior observations of Golgi reorganization in Arf1/COPI-depleted cells(23). Taken together, these findings suggest that the mechanism identified in this study has a broader role in regulating ER export of cargo proteins.

Cellular metabolites are essential for fulfilling the energy and biosynthetic demands of the cell. However, their potential involvement in regulating lysosomal enzyme trafficking remains poorly understood. While site-1 protease (S1P) is clearly required for lysosome biogenesis and function through proteolytic processing of GNPTAB, it remains unclear whether its role in lipid metabolic pathways also contributes to lysosomal regulation. Our study reveals a previously unrecognized link between lipid metabolism and lysosomal enzyme trafficking, suggesting that lipid metabolism may represent an additional regulatory layer that coordinates organelle function with cellular metabolic status. The biological significance and disease relevance of this regulation are worth further investigation.

Using proximity labeling, we identified a previously uncharacterized CTSL-interacting protein, Sccpdh2, as a novel regulator of CTSL transport. We propose that Sccpdh2 may function as a receptor mediating CTSL trafficking (SI Appendix, fig. S3I). One possibility is that Sccpdh2 acts as a cargo receptor for newly synthesized CTSL, facilitating its export from the ER to the Golgi. Retrieval of Sccpdh2 from the Golgi back to the ER would be required to sustain continuous CTSL export. Consistent with this hypothesis, our results indicate that bidirectional vesicle trafficking between the ER and the Golgi is critical for efficient CTSL transport. A key prediction of this model is that impaired retrieval of Sccpdh2 from the Golgi would deplete its ER pool, resulting in spatial separation between Sccpdh2 and CTSL. This is supported by the observations shown in Figure 4E. Alternatively, Sccpdh2 may function at a later stage, mediating CTSL transport from the Golgi to lysosomes. Further experiments will be required to distinguish between these possibilities. An additional important question is whether CTSL is the sole substrate of Sccpdh2 or whether Sccpdh2 serves as a broader receptor for multiple lysosomal or secretory proteins. Identifying additional substrates of Sccpdh2 will be essential for defining its general role in the secretory pathway and lysosomal biology.

Taken together, our findings uncover a previously unrecognized connection between lipid metabolism and lysosomal enzyme trafficking. These results expand the current understanding of lysosomal enzyme trafficking and provide a framework for future studies investigating how metabolic pathways coordinate with lysosomal function and cellular homeostasis.

## Online Methods

### Cell culture and transfection

*Drosophila* S2R+ cells were cultured with Schneider’s medium (Gibco) containing 10% heat-inactivated fetal bovine serum (Gibco) at 25 °C. Transfections of plasmids and dsRNAs into S2R+ cells were performed using Effectene (QIAGEN) following the manufacturer’s instructions. In brief, 3 × 10^6^ S2R+ cells were seeded into a single well of six-well plate prior to transfection. A total of 400 ng plasmid DNA or 20 μg dsRNA was diluted in Buffer EC to a final volume of 100 μL, then combined with 3.2 μL Enhancer and vortexed to generate a DNA-Enhancer mixture. Subsequently, 10 μL Effectene reagent was added to this mixture and vortexed again. After incubation at room temperature for 15 minutes to allow the transfection complexes to form, the mixture was added to the cells dropwise.

### Insertion of *GFP* into *CTSL* locus by homologous recombination

An sgRNA targeting the sequence TAGCGCCTCCAGCTATCCCC near the stop codon of *CTSL* was designed to induce a double-strand break. For the homologous recombination donor construct, ∼1 kb genomic regions upstream and downstream of the *CTSL* stop codon were cloned as homology arms flanking a GFP-T2A-Neo cassette. The sgRNA vector, a Cas9 expression vector, and the donor plasmid were co-transfected into *Drosophila* S2R+ cells. Transfected cells were subjected to neomycin selection for one month to establish a stable cell line.

### dsRNA synthesis and purification

dsRNAs were designed using DRSC/TRiP online tool Updated Targets of RNAi Reagents (UP-TORR) and synthesized according to DRSC/TRiP protocol. Briefly, T7 promoter sequence (TAATACGACTCACTATAGGG) were added to the 5’ end of target gene specific primers. dsRNA templates were amplified from genomic DNA by PCR. The resulting PCR amplicons were then used for in vitro transcription with the MEGAscript T7 Transcription Kit (Invitrogen) at 37 °C overnight to produce dsRNAs. Prior to transfection, dsRNAs were purified using the RNeasy Mini Kit (QIAGEN). All dsRNA primers used in this study are listed in the SI Appendix, Table S2.

### Western blotting

Cells were harvested and centrifuged to remove the culture medium, and the resulting cell pellet was washed once with PBS. Cells were then lysed in 1× cell lysis buffer (Cell Signaling Technology) supplemented with cOmplete™ Protease Inhibitor Cocktail (Roche). Protein concentrations were measured using a BCA protein assay kit (Thermo Fisher Scientific). The following antibodies were used for immunoblotting: anti-CTSL antibody (1:1000, MAB22591, R&D Systems), anti-Actin Rhodamine antibody (1:2500, AbD22606, Bio-Rad), anti-mouse IgG (H+L) secondary antibody, Alexa Fluor 800 (1:2500, Invitrogen, A32730). Western blotting images were taken by Bio-Rad ChemiDoc MP Imaging System.

### Fatty acid supplementation

Palmitate (10006627, Cayman Chemical Company) and myristate (13351, Cayman Chemical Company) were dissolved in ethanol and conjugated to fatty acid free bovine serum albumin (BSA) (A8806, Sigma-Aldrich). The fatty acids-BSA complexes were supplemented to S2R+ cells cultured with Schneider’s medium containing 10% heat-inactivated fetal bovine serum at 25 °C.

### Proximity labeling and pulldown assays

Plasmids encoding Bip-TurboID or CTSL-TurboID were transfected into S2R+ cells. 7 days after transfection, cells were incubated with 100 µM biotin for 2 hours to allow proximity-dependent biotinylation. Following biotin treatment, cells were washed four times with PBS and lysed in IP lysis buffer (Invitrogen) supplemented with cOmplete™ Protease Inhibitor Cocktail (Roche). Biotinylated proteins were subsequently enriched using streptavidin magnetic beads and analyzed by mass spectrometry.

### Recombinant protein expression and purification

The N-terminal region of *Sccpdh2* was cloned into the pET-26b vector. The resulting plasmid was transformed into BL21 bacterial cells, and recombinant protein expression was induced with 0.3 mM IPTG at 15 °C overnight. Following induction, bacteria were collected by centrifugation at 6000 × g for 15 min at 4 °C. The periplasmic fraction was extracted using B-PER II reagent (Thermo Fisher Scientific, 78260). The periplasmic extract was passed through a 0.22 µm filter, supplemented with 10 mM imidazole, and subjected to purification by Ni²⁺ affinity chromatography (Cytiva, 17531801). The sample was loaded onto a gravity-flow column containing 1 mL Ni²⁺ resin that had been pre-equilibrated with TBS containing 10 mM imidazole. The resin was washed three times with 20 mL TBS supplemented with 20 mM imidazole. Bound nanobody-ALFA-His protein was eluted in 1 mL fractions using TBS containing stepwise increases in imidazole concentration (100 mM, 250 mM, and 500 mM). The eluted protein samples were analyzed by stain-free protein gels (Bio-Rad) and stored at 4 °C.

## Acknowledgments

We thank John Asara at the BIDMC-Harvard mass spectrometry facility for the mass spectrometry analysis of proximity labeling samples. This work is supported by NIH/NIAMS 5R01AR057352 (N.P.) and the HMS-SNUH Research and Education Collaboration program. N.P. is an investigator of Howard Hughes Medical Institute. This article is subject to HHMI’s Immediate Access to Research policy, which requires that this article be made publicly available as initial and revised preprints deposited on a designated preprint server under a CC BY 4.0 license.

