## Supplementary Information for "Lipids Regulate Export of Lysosomal Enzymes from the Endoplasmic Reticulum"

PNAS

### Supporting Information for
Lipids Regulate Export of Lysosomal Enzymes from the Endoplasmic Reticulum

Baolong Xia, Myeonghoon Han, Isaac Park, Norbert Perrimon

Corresponding author: Norbert Perrimon


**This PDF file includes:**

Figures S1 to S3
Tables S1 to S2

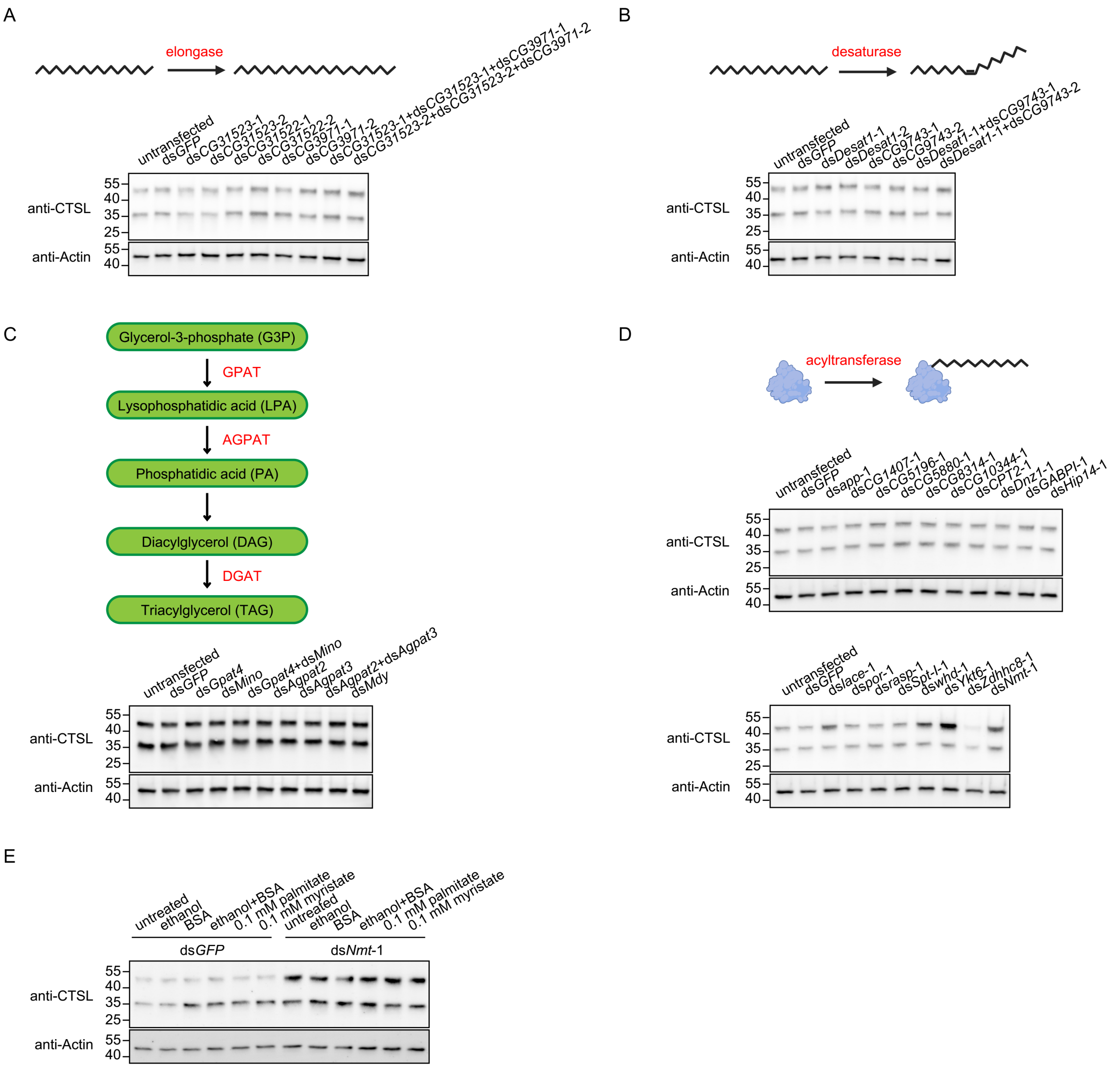


**Fig. S1. Fatty acid elongation, desaturation, and triglyceride synthesis are dispensable for CTSL trafficking.**

A. Western blot analysis of CTSL in cells transfected with dsRNAs targeting elongases expressed in Drosophila S2R+ cells.

B. Western blot analysis of CTSL in cells transfected with dsRNAs targeting desaturases expressed in Drosophila S2R+ cells.

C. Western blot analysis of CTSL in cells transfected with dsRNAs targeting enzymes involved in triglyceride synthesis.

D. Western blot analysis of CTSL in cells transfected with dsRNAs targeting acyltransferase involved in protein acylation.

E. Western blot analysis of CTSL in Nmt knockdown cells supplemented with 0.1 mM palmitate (C16:0) or myristate (C14:0).

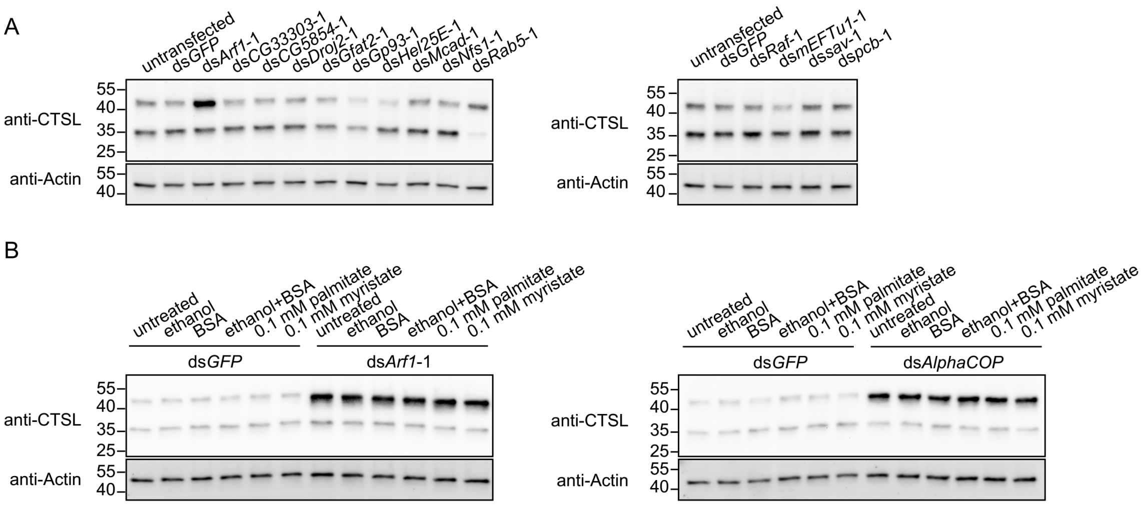


**Supplementary Figure 2. Arf1/COPI regulates CTSL transport independently of fatty acids.**

A. Western blot analysis of CTSL in cells following depletion of known CTSL-interacting proteins.

B. Western blot analysis of CTSL in Arf1 or AlphaCOP knockdown cells supplemented with 0.1 mM palmitate (C16:0) or myristate (C14:0).

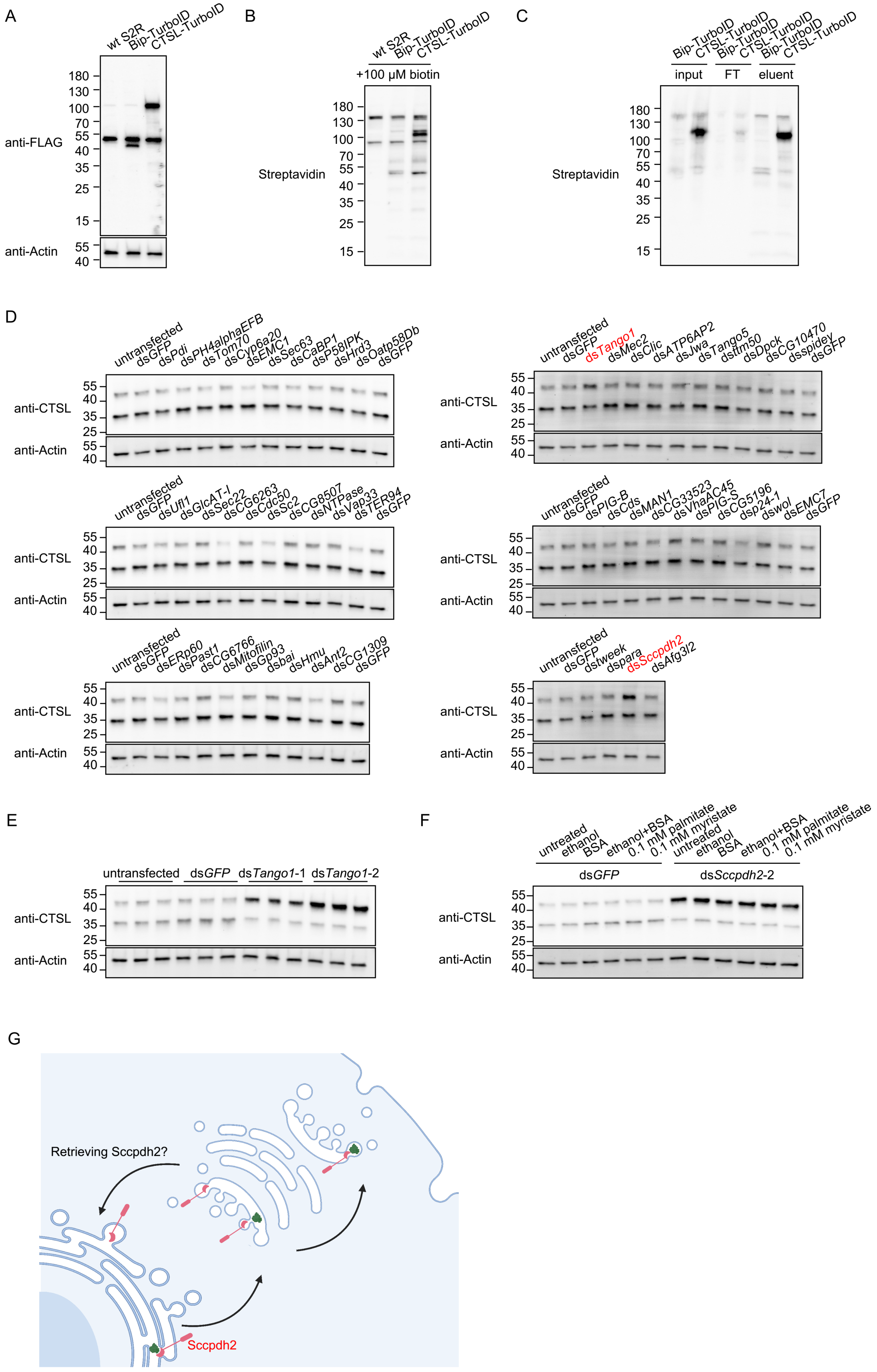
A
wt S2R
Bip-TurboID
CTSL-TurboID
anti-FLAG
anti-Actin
180
130
100
70
55
40
35
25
15
B
+100 μM biotin
Streptavidin
C
input
FT
eluent
D
anti-CTSL
untransfected
dsGFP
dsPdi
dsPH4alphaEFB
dsTom70
dsCyp6a20
dsEMC1
dsSec63
dsCaBP1
dsP58IPK
dsHrd3
dsOatp58Db
dsTango1
dsMec2
dsClic
dsATP6AP2
dsJwa
dsTango5
dsttm50
dsDpck
dsCG10470
dsspidey
dsUfl1
dsGlcAT-I
dsSec22
dsCG6263
dsCdc50
dsSc2
dsCG8507
dsNTPase
dsVap33
dsTER94
dsPIG-B
dsCds
dsMAN1
dsCG33523
dsVhaAC45
dsPIG-S
dsCG5196
dsp24-1
dswol
dsEMC7
dsERp60
dsPast1
dsCG6766
dsMitofilin
dsGp93
dsbai
dsHmu
dsAnt2
dsCG1309
dstweek
dspara
dsSccpdh2
dsAfg3l2
E
dsTango1-1
dsTango1-2
F
untreated
ethanol
BSA
ethanol+BSA
0.1 mM palmitate
0.1 mM myristate
dsSccpdh2-2
G
Retrieving Sccpdh2?
Sccpdh2

**Supplementary Figure 3. Proximity labeling identifies novel regulators of CTSL transport.**

A. Western blot analysis of FLAG-tagged BiP-TurboID and CTSL-TurboID expression.

B. Western blot analysis of biotinylated proteins in cells transfected with plasmids encoding BiP-TurboID and CTSL-TurboID. Cells were treated with 100 μM biotin for 2 hours to allow proximity-dependent biotinylation.

C. Western blot analysis of biotinylated proteins in the pulldown samples. FT, flow-through.

D. Western blot analysis of CTSL in following knockdown of candidate genes identified by proximity labeling.

E. Western blot analysis of CTSL in Tango1 knockdown cells. Two different non-overlapping dsRNAs targeting Tango1 were used.

F. Western blot analysis of CTSL in Sccpdh2 knockdown cells supplemented with 0.1 mM palmitate (C16:0) or myristate (C14:0).

G. Western blot analysis of ImpL2 in cell lysate following knockdown of SREBP, Arf1, or Sccpdh2. The red arrowhead indicates the ImpL2 bands and the asterisk indicates the nonspecific bands.

H. Immunostaining of Golgin84 in dsGFP and dsSREBP transfected cells.

I. Model: Sccpdh2 functions as a cargo receptor to facilitate CTSL export from the ER to the Golgi, and its retrieval is required to sustain continuous CTSL trafficking.

**Table S1. Mass spectrometry data of proximity labeling samples.**

| Accession Number | counts in BiP-TurboID | counts in CTSL-TurboID |
|---|---|---|
| Q99323 | 747 | 841 |
| Q0E9E2 | 602 | 887 |
| A1Z784 | 287 | 848 |
| P10987 | 387 | 295 |
| P02572 | 382 | 293 |
| P11147 | 245 | 391 |
| P02828 | 224 | 317 |
| P02574 | 208 | 215 |
| Q9VAY2 | 79 | 309 |
| Q9V9T5 | 100 | 254 |
| P29844 | 100 | 220 |
| Q24560 | 92 | 190 |
| P08736 | 127 | 117 |
| P06603 (+1) | 79 | 122 |
| P08841 | 63 | 132 |
| P29845 | 62 | 131 |
| Q9NJH0 | 65 | 111 |
| Q94920 | 89 | 86 |
| A1Z6H7 | 65 | 103 |
| Q26365 | 62 | 105 |
| O97125 | 83 | 57 |
| P35381 | 46 | 95 |
| O46067 | 34 | 103 |
| Q8MST5 | 35 | 100 |
| A1Z843 | 49 | 83 |
| P02825 (+1) | 69 | 59 |
| Q05825 | 35 | 92 |
| Q8INI8 (+3) | 69 | 56 |
| Q9VN21 | 66 | 56 |
| P21187 | 41 | 73 |
| Q9W0S7 | 41 | 70 |
| Q9VV75 | 43 | 61 |
| Q9W392 | 31 | 73 |
| Q95TL8 | 28 | 74 |
| P19109 | 19 | 81 |
| Q9VEX6 | 44 | 56 |
| Q3YMU0 | 17 | 82 |
| Q9VHL2 | 48 | 51 |
| O62619 | 26 | 68 |

| | | |
|---|---|---|
| P24156 | 43 | 51 |
| Q06559 | 40 | 54 |
| Q9VXQ5 | 30 | 64 |
| Q02748 | 35 | 55 |
| Q9VZ23 | 31 | 58 |
| Q7K3J0 | 19 | 68 |
| A1Z6L9 | 25 | 61 |
| A8DYI6 | 36 | 49 |
| P54357 | 49 | 34 |
| A1Z729 | 28 | 54 |
| Q8T0L3 | 24 | 57 |
| Q9VRL8 | 34 | 47 |
| P55830 | 29 | 50 |
| Q95028 | 28 | 49 |
| P48605 | 37 | 39 |
| P29742 | 32 | 43 |
| P45594 | 39 | 36 |
| P09180 | 26 | 47 |
| Q8MLY8 | 23 | 50 |
| Q9V9Q4 | 34 | 34 |
| Q01604 | 16 | 51 |
| Q9VK59 | 33 | 32 |
| P13469 | 29 | 35 |
| C0HKA0 (+1) | 25 | 37 |
| P02515 | 35 | 26 |
| Q9VLC5 | 7 | 54 |
| B7Z001 | 10 | 49 |
| O16797 | 25 | 33 |
| Q27331 | 15 | 43 |
| Q9VHP0 | 26 | 32 |
| P41042 | 16 | 41 |
| P54399 | 1 | 56 |
| P31009 | 23 | 32 |
| Q9VN25 | 12 | 43 |
| Q9VK69 | 12 | 42 |
| P22700 | 20 | 33 |
| P23226 | 19 | 34 |
| Q95U38 | 15 | 38 |
| Q9VB10 | 24 | 29 |
| O18332 | 25 | 27 |
| P07909 | 40 | 12 |
| P48810 | 26 | 24 |
| A8DZ29 | 16 | 35 |

| | | |
|---|---|---|
| Q7KKI0 | 20 | 30 |
| Q9VFF0 | 16 | 33 |
| A1Z9E3 | 10 | 38 |
| P02516 | 16 | 32 |
| P55841 | 15 | 33 |
| Q08473 | 30 | 18 |
| Q8IQZ7 | 19 | 29 |
| O62526 | 18 | 29 |
| O01666 | 17 | 29 |
| P41093 | 19 | 27 |
| P54397 | 19 | 27 |
| Q9VNE9 | 18 | 28 |
| Q7KN75 | 18 | 27 |
| A0A0B4K7J2 | 18 | 27 |
| P12613 | 16 | 29 |
| P41374 | 21 | 23 |
| Q9W237 | 11 | 33 |
| P18101 | 20 | 23 |
| P55935 | 14 | 29 |
| Q9V470 | 8 | 34 |
| Q9VS34 | 11 | 31 |
| Q9VH64 | 6 | 36 |
| Q7JYX2 | 15 | 27 |
| O96827 | 27 | 14 |
| P48149 | 15 | 26 |
| Q24208 | 24 | 17 |
| Q9V9A7 | 7 | 34 |
| Q9VHC7 | 26 | 15 |
| Q9W327 | 12 | 28 |
| Q9W0P5 | 11 | 29 |
| Q9W1H8 | 20 | 20 |
| Q9V3A8 | 12 | 27 |
| Q9VVU2 | 13 | 26 |
| Q9VKB0 | 15 | 23 |
| P02518 | 17 | 21 |
| P13060 | 18 | 20 |
| Q07327 | 18 | 20 |
| Q7K0E6 | 12 | 26 |
| Q9W5R8 | 9 | 29 |
| O18640 | 14 | 23 |
| Q9VIE8 | 6 | 30 |
| Q9W0B8 | 6 | 30 |
| P32100 | 6 | 29 |

| | | |
|---|---|---|
| P20432 | 15 | 19 |
| P29327 | 10 | 24 |
| P38979 | 15 | 19 |
| P46863 | 14 | 20 |
| Q7KMQ0 | 15 | 19 |
| Q8T4G5 | 9 | 25 |
| P41094 | 14 | 19 |
| P50887 | 14 | 19 |
| Q9VKD3 | 6 | 27 |
| A1ZAX1 | 9 | 23 |
| P36179 | 12 | 20 |
| P48809 | 11 | 21 |
| Q9VKW5 | 6 | 26 |
| Q27268 | 7 | 24 |
| Q76NQ0 | 8 | 23 |
| Q7K110 | 12 | 19 |
| Q9V3Q4 | 7 | 24 |
| P61209 | 9 | 22 |
| Q7KN94 | 7 | 23 |
| Q9VWG3 | 4 | 26 |
| P46222 | 13 | 17 |
| P52295 | 9 | 20 |
| P07486 | 12 | 17 |
| P25161 | 18 | 11 |
| P48159 | 11 | 18 |
| Q9VBV5 | 15 | 14 |
| X2JC79 | 10 | 19 |
| Q94522 | 3 | 24 |
| Q07171 | 16 | 12 |
| Q9W0A8 | 9 | 19 |
| Q9VQL1 | 5 | 22 |
| Q02645 | 8 | 19 |
| Q9V9W3 | 4 | 23 |
| Q9W1V3 | 10 | 17 |
| Q9W3Y3 | 8 | 19 |
| Q9XZJ4 | 10 | 17 |
| Q9V438 | 2 | 25 |
| Q9VW54 | 3 | 24 |
| P28668 | 7 | 19 |
| P84040 | 10 | 16 |
| Q9V597 | 13 | 13 |
| Q9VWI2 | 12 | 14 |
| Q9VZL1 | 21 | 5 |

| | | |
|---|---|---|
| P41092 | 7 | 19 |
| Q7KN62 | 4 | 22 |
| P50882 | 4 | 22 |
| Q9VZS5 | 8 | 17 |
| Q9VGA0 | 10 | 15 |
| Q9VSD6 | 15 | 10 |
| Q9W4M9 | 6 | 19 |
| Q9VN44 | 4 | 21 |
| Q9VHA8 | 2 | 23 |
| O17445 | 9 | 15 |
| Q24319 | 7 | 17 |
| Q94518 | 14 | 10 |
| Q9V447 | 18 | 6 |
| Q9VKZ8 | 10 | 14 |
| Q7KR10 | 4 | 20 |
| Q7KB18 | 22 | 2 |
| O18413 | 6 | 17 |
| P29310 | 14 | 9 |
| Q03334 | 5 | 18 |
| Q24492 | 11 | 12 |
| Q7K3D4 | 11 | 12 |
| Q9VHS8 | 19 | 4 |
| Q7K2G1 | 13 | 9 |
| Q8I0G5 | 9 | 13 |
| Q9V3G1 | 10 | 12 |
| Q9VH95 | 6 | 16 |
| Q9VEZ3 | 6 | 16 |
| Q04499 | 5 | 17 |
| Q9VUQ5 | 5 | 17 |
| Q8T9B6 | 8 | 14 |
| O18333 | 6 | 16 |
| M9NF14 | 1 | 21 |
| Q0E8E8 | 7 | 14 |
| Q9VT61 | 5 | 16 |
| Q9W2N0 | 12 | 9 |
| P13607 | 6 | 15 |
| Q7KUT2 | 5 | 15 |
| O18404 | 4 | 16 |
| Q8SWU7 | 8 | 12 |
| P22464 | 8 | 12 |
| P02517 | 4 | 16 |
| Q9V455 | 7 | 13 |
| Q9V434 | 4 | 16 |

| | | |
|---|---|---|
| Q9VB46 | 4 | 16 |
| A1ZBE9 | 5 | 15 |
| A1Z9N0 | 5 | 14 |
| Q9V9M7 | 7 | 12 |
| Q9VY91 | 9 | 10 |
| Q9V3P6 | 4 | 15 |
| Q7KLV9 | 4 | 15 |
| O18334 | 6 | 13 |
| Q9TVM2 | 7 | 12 |
| Q9VTU4 | 4 | 15 |
| P91926 | 5 | 14 |
| P55828 | 4 | 15 |
| Q9VA69 | 0 | 19 |
| P45437 | 5 | 13 |
| P48603 | 8 | 10 |
| Q95029 | 4 | 14 |
| Q7KMP8 | 8 | 10 |
| Q7JVI3 | 5 | 13 |
| Q0KHZ6 | 3 | 15 |
| A1Z934 | 6 | 12 |
| Q01637 | 3 | 15 |
| Q7KMJ6 | 2 | 16 |
| Q8MZI3 | 3 | 15 |
| E1JGL8 | 7 | 10 |
| O61231 | 7 | 10 |
| P32234 | 4 | 13 |
| Q9VAW5 | 7 | 10 |
| Q9W3W8 | 5 | 12 |
| Q9VV60 | 4 | 13 |
| Q9VJ86 | 4 | 13 |
| P31409 | 4 | 13 |
| Q9VRP3 | 5 | 12 |
| Q8IPX7 | 9 | 8 |
| P08928 | 6 | 11 |
| P46223 | 3 | 14 |
| P35600 | 3 | 13 |
| P17704 | 5 | 11 |
| P48601 | 5 | 11 |
| Q26454 | 6 | 10 |
| Q95RI5 | 8 | 8 |
| Q9V3V6 | 8 | 8 |
| Q9VJY6 | 4 | 12 |
| A1ZBW0 | 10 | 6 |

| | | |
|---|---|---|
| Q9VSS2 | 8 | 8 |
| P14199 | 9 | 7 |
| P52304 | 7 | 9 |
| Q9VUZ0 | 7 | 9 |
| Q9W414 | 3 | 13 |
| M9PBV2 | 2 | 14 |
| Q8SWR8 | 5 | 11 |
| Q8T079 | 4 | 12 |
| Q9VBN5 | 2 | 14 |
| Q9V7D2 | 1 | 15 |
| Q95WY3 | 3 | 12 |
| A1ZAB5 | 7 | 8 |
| Q9V3J1 | 5 | 10 |
| Q9VH81 | 8 | 7 |
| Q9VAL7 | 4 | 11 |
| O02195 | 7 | 8 |
| Q9VPR1 | 3 | 12 |
| A0A0B4KFE4 | 5 | 10 |
| Q9XZU1 | 5 | 10 |
| Q9VVA7 | 4 | 11 |
| Q9Y105 | 3 | 12 |
| P41375 | 3 | 12 |
| Q9VMA7 | 3 | 12 |
| Q9W4N8 | 2 | 13 |
| Q9VSA3 | 1 | 14 |
| O15971 | 4 | 11 |
| Q9VYW4 | 4 | 9 |
| Q9V436 | 4 | 10 |
| P41572 | 4 | 10 |
| Q8IQV9 | 3 | 11 |
| Q9VGP4 | 5 | 9 |
| Q9VNB9 | 5 | 9 |
| O76742 | 6 | 8 |
| Q9VPQ7 | 9 | 5 |
| Q9VWH4 | 3 | 11 |
| P40945 | 0 | 13 |
| A8JNJ6 | 4 | 9 |
| A8JNK1 | 6 | 7 |
| Q9VF03 | 7 | 6 |
| Q9W2E7 | 9 | 4 |
| Q9VRL1 | 11 | 2 |
| Q9Y134 | 3 | 10 |
| Q9NFU0 | 6 | 7 |

| | | |
|---|---|---|
| Q9GU68 | 4 | 9 |
| Q7K2L7 | 5 | 8 |
| Q9VKY2 | 2 | 11 |
| Q94524 | 3 | 10 |
| Q9VKC8 | 0 | 13 |
| Q9W5X0 | 0 | 10 |
| P48591 | 5 | 7 |
| Q9VA91 | 7 | 5 |
| Q9W229 | 6 | 6 |
| Q9Y162 | 5 | 7 |
| Q7KSM5 | 6 | 6 |
| Q9V3I2 | 5 | 7 |
| Q9VKM7 | 4 | 8 |
| Q9VFT4 | 5 | 7 |
| P45889 | 4 | 8 |
| Q9VBX1 | 6 | 6 |
| Q9VD58 | 7 | 5 |
| Q8IRD3 | 3 | 9 |
| P05205 | 7 | 5 |
| Q9VQ29 | 4 | 8 |
| P91891 | 4 | 8 |
| Q9V3J4 | 5 | 7 |
| P40320 | 4 | 8 |
| P17210 | 3 | 9 |
| P15348 | 2 | 10 |
| Q9W3Q0 | 4 | 8 |
| Q7K4L8 | 2 | 10 |
| Q9V6U9 | 1 | 11 |
| P02283 | 3 | 8 |
| Q9VQG4 | 6 | 5 |
| P37276 | 3 | 8 |
| Q94517 | 6 | 5 |
| Q9VQR8 | 4 | 7 |
| Q9W3K5 | 3 | 8 |
| Q7KSQ0 | 4 | 7 |
| A0A140SRF8 | 8 | 3 |
| P36241 | 4 | 7 |
| Q0E9B6 | 7 | 4 |
| Q9VDK7 | 4 | 7 |
| Q9V3Z4 | 5 | 6 |
| Q9VQX4 | 3 | 8 |
| Q9V3D9 | 4 | 7 |
| P04359 | 3 | 8 |

| | | |
|---|---|---|
| Q7KM15 | 2 | 9 |
| A1ZB71 | 3 | 8 |
| Q8IRG6 | 3 | 8 |
| Q9V461 | 8 | 3 |
| Q24133 | 3 | 8 |
| Q27415 | 2 | 9 |
| Q1RL13 | 3 | 8 |
| Q8MSW0 | 3 | 8 |
| Q9VD22 | 2 | 9 |
| Q24253 | 1 | 10 |
| Q9VZL3 | 1 | 10 |
| Q9VPY2 | 7 | 3 |
| P49071 | 6 | 4 |
| Q7K4Q5 | 3 | 7 |
| Q8SX76 | 5 | 5 |
| Q9VQX6 | 3 | 7 |
| Q9VAI4 | 6 | 4 |
| A1Z8U0 | 4 | 6 |
| Q9VVU6 | 6 | 4 |
| Q7JZW2 | 7 | 3 |
| Q9VXG4 | 2 | 8 |
| Q9VNE2 | 3 | 7 |
| Q9VJZ1 | 3 | 7 |
| P35128 | 3 | 7 |
| Q9VPJ0 | 4 | 6 |
| O76511 | 6 | 4 |
| P55035 | 5 | 5 |
| P08985 | 4 | 6 |
| Q8SY19 | 6 | 4 |
| P53034 | 2 | 8 |
| P08181 | 3 | 7 |
| Q9VT75 | 6 | 4 |
| Q7KW39 | 3 | 7 |
| Q9VVW7 | 4 | 6 |
| P49630 | 1 | 9 |
| Q9VVA6 | 3 | 7 |
| P29413 | 2 | 8 |
| Q9VM50 | 5 | 5 |
| Q7KLX3 | 2 | 8 |
| O77410 | 3 | 7 |
| Q7KVX1 | 3 | 7 |
| M9PC99 | 0 | 10 |
| Q9V773 | 0 | 10 |

| | | |
|---|---|---|
| Q9VHY6 | 0 | 10 |
| Q7KT60 | 2 | 6 |
| Q9VE85 | 2 | 7 |
| A0A0B4KI34 | 4 | 5 |
| A0A0B4LGM5 | 5 | 4 |
| Q9VLT5 | 5 | 4 |
| Q24478 | 5 | 4 |
| Q9VUV9 | 3 | 6 |
| Q9VZ20 | 5 | 4 |
| P08120 | 3 | 6 |
| Q9W2D9 | 3 | 6 |
| Q9VTZ4 | 3 | 6 |
| Q9W555 | 2 | 7 |
| Q9W2I2 | 4 | 5 |
| P40417 | 2 | 7 |
| Q7KRW8 | 3 | 6 |
| Q960W6 | 5 | 4 |
| Q27237 | 1 | 8 |
| Q9VZS3 | 7 | 2 |
| Q9V405 | 2 | 7 |
| Q0KI98 | 1 | 8 |
| Q8IPV3 | 4 | 3 |
| A0A0B4KHI4 | 1 | 7 |
| Q24368 | 1 | 7 |
| P41126 | 4 | 4 |
| Q86BI3 | 5 | 3 |
| Q9W0M4 | 4 | 4 |
| Q7KUD4 | 3 | 5 |
| Q9V9S8 | 5 | 3 |
| Q9VPM7 | 2 | 6 |
| Q04047 | 2 | 6 |
| P52034 | 5 | 3 |
| Q9VIU7 | 2 | 6 |
| Q9VP77 | 3 | 5 |
| O02649 | 2 | 6 |
| Q9VP57 | 4 | 4 |
| Q9VZI1 | 4 | 4 |
| Q9VN50 | 3 | 5 |
| P23128 | 2 | 6 |
| Q05344 | 6 | 2 |
| Q8MT18 | 5 | 3 |
| Q9U9Q1 | 2 | 6 |
| Q9U9Q4 | 3 | 5 |

| | | |
|---|---|---|
| Q8MT58 | 3 | 5 |
| Q7K0D8 | 4 | 4 |
| Q9W1G7 | 2 | 6 |
| P25843 | 1 | 7 |
| P00522 | 2 | 6 |
| Q8T8W3 | 1 | 7 |
| P13395 | 2 | 6 |
| C0HL66 (+2) | 6 | 2 |
| A0A0B4LID7 | 1 | 7 |
| Q94901 | 2 | 6 |
| Q95PE4 | 6 | 2 |
| Q9NCC3 | 1 | 7 |
| Q9VKC7 | 1 | 7 |
| Q9V460 | 1 | 7 |
| Q7K5K3 | 1 | 7 |
| Q9VW59 | 1 | 7 |
| P91928 | 1 | 7 |
| M9PFR8 (+1) | 0 | 8 |
| O62621 | 0 | 8 |
| Q9VRL0 | 0 | 8 |
| Q9VZQ1 | 4 | 3 |
| P20480 | 2 | 5 |
| Q9V4P1 | 3 | 4 |
| Q86S05 | 3 | 4 |
| Q9VK25 | 4 | 3 |
| Q9VAJ9 | 3 | 4 |
| Q9VD14 | 3 | 4 |
| Q9W3C1 | 2 | 5 |
| Q9VJ19 | 2 | 5 |
| P91875 | 2 | 5 |
| Q9W2X6 | 3 | 4 |
| A0A0B4KF06 | 2 | 5 |
| O18388 | 1 | 6 |
| Q23979 | 1 | 6 |
| Q9VIF2 | 3 | 4 |
| Q7K4Y0 | 1 | 6 |
| Q9VN86 | 3 | 4 |
| Q9VSU6 | 2 | 5 |
| Q86B44 | 4 | 3 |
| Q9VLU0 | 3 | 4 |
| Q7KRY6 | 2 | 5 |
| Q9VM14 | 1 | 6 |
| Q9Y0Y5 | 2 | 5 |

| | | |
|---|---|---|
| Q9VEK8 | 2 | 5 |
| Q9W334 | 3 | 4 |
| A1Z6H6 | 1 | 6 |
| Q9VCK0 | 2 | 5 |
| Q9V397 | 1 | 6 |
| Q8SXY6 | 1 | 6 |
| Q9VVL8 | 2 | 5 |
| Q9W253 | 2 | 5 |
| M9PGG8 | 7 | 0 |
| Q7KVQ0 | 1 | 6 |
| Q8IRE4 | 0 | 7 |
| Q9VS57 | 0 | 7 |
| Q8STG9 | 0 | 7 |
| Q7KRU8 | 0 | 7 |
| Q9VCX7 | 2 | 3 |
| M9MRD1 | 3 | 3 |
| Q95U54 | 2 | 4 |
| Q9W350 | 2 | 4 |
| Q9W5E4 | 4 | 2 |
| Q9W252 | 3 | 3 |
| Q9I7T7 | 3 | 3 |
| Q7K012 | 3 | 3 |
| Q9VSK9 | 2 | 4 |
| Q9VB22 | 2 | 4 |
| Q9VAN7 | 2 | 4 |
| Q9NHD5 | 2 | 4 |
| M9PI41 | 2 | 4 |
| Q9VW52 | 2 | 4 |
| Q9VC57 | 3 | 3 |
| Q9V3K9 | 2 | 4 |
| Q86BS3 | 4 | 2 |
| Q9VRG8 | 2 | 4 |
| Q9VGA4 | 3 | 3 |
| Q9NBD7 | 2 | 4 |
| Q8MLV1 | 3 | 3 |
| A1ZA77 | 3 | 3 |
| P23257 | 2 | 4 |
| Q9V5C6 | 4 | 2 |
| Q9VZF1 | 3 | 3 |
| O77263 | 2 | 4 |
| Q9VWL0 | 1 | 5 |
| P54622 | 2 | 4 |
| Q9VGW7 | 3 | 3 |

| | | |
|---|---|---|
| Q9VH78 | 3 | 3 |
| Q9VHE5 | 1 | 5 |
| Q9V3W1 | 2 | 4 |
| P19889 | 3 | 3 |
| Q8T3U2 | 2 | 4 |
| Q9VK44 | 2 | 4 |
| P08111 | 2 | 4 |
| A1ZAW0 | 2 | 4 |
| Q9VUR3 | 2 | 4 |
| P24785 | 1 | 5 |
| P08645 | 1 | 5 |
| Q9VY78 | 1 | 5 |
| Q9V431 | 1 | 5 |
| Q9VAC4 | 1 | 5 |
| Q9VGQ1 | 3 | 3 |
| P12881 | 2 | 4 |
| P48598 | 2 | 4 |
| A1Z8S6 | 2 | 4 |
| Q9VKW3 | 1 | 5 |
| Q7K2B0 | 1 | 5 |
| A1Z6P3 | 1 | 5 |
| Q9VNH2 | 3 | 3 |
| Q94511 | 1 | 5 |
| Q9VUK8 | 1 | 5 |
| Q9V3W2 | 2 | 4 |
| Q9W141 | 1 | 5 |
| Q9VPR3 | 1 | 5 |
| Q24186 | 1 | 5 |
| Q9XZ06 | 1 | 5 |
| Q9VZE4 | 1 | 5 |
| Q9VXN4 | 1 | 5 |
| Q9VVU1 | 0 | 6 |
| Q9VST4 | 0 | 6 |
| Q94883 | 0 | 6 |
| Q9VW13 | 0 | 6 |
| Q9VN95 | 0 | 6 |
| Q7PLI7 | 0 | 6 |
| Q9VXF9 | 1 | 3 |
| P26270 | 1 | 4 |
| Q8SXP0 | 1 | 4 |
| Q9VLX0 | 2 | 3 |
| Q8I941 | 1 | 3 |
| P25171 | 3 | 2 |

| | | |
|---|---|---|
| Q9VJD1 | 1 | 4 |
| Q9V9R2 | 1 | 4 |
| Q9U4L6 | 2 | 3 |
| Q9VC05 | 2 | 3 |
| Q9VYQ8 | 2 | 3 |
| Q9VRP5 | 2 | 3 |
| Q9W0C5 | 2 | 3 |
| Q9W074 | 2 | 3 |
| Q8IRH5 | 2 | 3 |
| A1ZBL9 | 2 | 3 |
| Q9VTX8 | 2 | 3 |
| Q9V463 | 3 | 2 |
| Q9VKB2 | 2 | 3 |
| Q7K1Q7 | 2 | 3 |
| O62530 | 3 | 2 |
| Q9VVV7 | 2 | 3 |
| Q9VQQ0 | 2 | 3 |
| Q9W002 | 2 | 3 |
| Q9VND3 | 2 | 3 |
| P32392 | 3 | 2 |
| Q9VXR5 | 2 | 3 |
| Q7JR58 | 1 | 4 |
| Q9VIP0 | 2 | 3 |
| Q9VVL7 | 2 | 3 |
| Q9V3E9 | 1 | 4 |
| Q8INM3 | 1 | 4 |
| Q8MRM0 | 1 | 4 |
| Q9VQG1 | 1 | 4 |
| Q9VZP5 | 1 | 4 |
| Q9W1H5 | 1 | 4 |
| Q94533 | 2 | 3 |
| Q9VII9 | 1 | 4 |
| O61613 | 1 | 4 |
| Q9W0G1 | 4 | 1 |
| P40796 | 4 | 1 |
| Q9VKJ6 | 4 | 1 |
| Q9VIH1 | 2 | 3 |
| O77277 | 1 | 4 |
| Q9VKW1 | 2 | 3 |
| Q7K4C7 | 2 | 3 |
| Q9VI58 | 1 | 4 |
| Q9VGK7 | 1 | 4 |
| Q9VHG4 | 1 | 4 |

| | | |
|---|---|---|
| Q9VKJ3 | 1 | 4 |
| Q9V9K7 | 1 | 4 |
| Q05913 | 1 | 4 |
| Q9VEN3 | 2 | 3 |
| Q9W179 | 2 | 3 |
| Q24179 | 2 | 3 |
| Q9VEQ2 | 1 | 4 |
| Q0KIB3 | 1 | 4 |
| Q8T3L6 | 1 | 4 |
| Q9V9X4 | 1 | 4 |
| Q24439 | 3 | 2 |
| P13008 | 2 | 3 |
| Q9VQQ6 | 0 | 5 |
| Q5BI50 | 1 | 4 |
| P04052 | 0 | 5 |
| Q9V3W7 | 1 | 4 |
| Q9VDV2 | 0 | 5 |
| E1JIH4 | 0 | 5 |
| Q9VQB4 | 0 | 5 |
| O61444 | 0 | 5 |
| A8JNU6 | 0 | 5 |
| P23572 | 0 | 5 |
| Q9V415 | 0 | 5 |
| Q9VCE0 | 0 | 5 |
| Q9VM69 | 0 | 5 |
| Q9VRV8 | 0 | 5 |
| Q9V3G7 | 0 | 5 |
| Q9VID7 | 0 | 5 |
| Q9XYU0 | 0 | 5 |
| Q9VBU9 | 0 | 5 |
| Q9W270 | 0 | 5 |
| Q8MKN0 | 1 | 3 |
| A0A0B4K765 | 1 | 3 |
| X2JAM4 | 2 | 2 |
| Q9VXY3 | 2 | 2 |
| Q9VLT1 | 2 | 2 |
| P00967 | 2 | 2 |
| Q9VMX1 | 3 | 1 |
| Q9VND8 | 1 | 3 |
| Q9VFE7 | 2 | 2 |
| Q9VVH5 | 2 | 2 |
| Q9VQR9 | 2 | 2 |
| O62602 | 1 | 3 |

| | | |
|---|---|---|
| P39018 | 1 | 3 |
| O77086 | 2 | 2 |
| Q9VQI5 | 1 | 3 |
| Q9W1Q8 | 2 | 2 |
| M9PHV4 | 1 | 3 |
| Q8SXU3 | 1 | 3 |
| Q9VKZ7 | 1 | 3 |
| Q9V3V0 | 2 | 2 |
| Q9VKC1 | 1 | 3 |
| A1Z9K0 | 1 | 3 |
| Q9VIE7 | 1 | 3 |
| Q7KHK9 | 2 | 2 |
| Q9VYA7 | 2 | 2 |
| A0A0B4KEJ7 | 3 | 1 |
| Q7K4I5 | 3 | 1 |
| Q9VUH8 | 2 | 2 |
| O18335 | 2 | 2 |
| M9NF32 | 1 | 3 |
| Q9VLB7 | 1 | 3 |
| O76902 | 2 | 2 |
| Q9VGF1 | 1 | 3 |
| A0A0B4KGS4 | 2 | 2 |
| P56175 | 2 | 2 |
| Q9W0S9 | 2 | 2 |
| Q9Y109 | 2 | 2 |
| P20477 | 1 | 3 |
| A1ZB68 | 1 | 3 |
| Q9VLK2 | 1 | 3 |
| P51406 | 3 | 1 |
| Q95U34 | 1 | 3 |
| Q9V345 | 1 | 3 |
| Q0KHQ1 | 1 | 3 |
| Q9XZ61 | 1 | 3 |
| Q9VQ78 | 1 | 3 |
| Q960Z0 | 1 | 3 |
| D5AEK7 | 1 | 3 |
| Q9VL70 | 1 | 3 |
| Q6NN55 | 1 | 3 |
| P20354 | 1 | 3 |
| M9PDW8 | 0 | 3 |
| Q9VPH2 | 1 | 3 |
| Q9VWL7 | 3 | 1 |
| Q9VHX2 | 1 | 3 |

| | | |
|---|---|---|
| P22817 | 3 | 1 |
| A1Z9M5 | 0 | 3 |
| Q9VQE0 | 1 | 3 |
| Q6IDD9 | 0 | 4 |
| Q9VA73 | 1 | 3 |
| Q7JZR5 | 2 | 2 |
| A0A0B4LEZ3 | 1 | 3 |
| Q9VQ79 | 2 | 2 |
| P55824 | 1 | 3 |
| Q9VRJ4 | 1 | 3 |
| Q9V7N5 | 1 | 3 |
| Q9VD51 | 1 | 3 |
| Q9VXE6 | 0 | 4 |
| Q9VJZ7 | 1 | 3 |
| Q9V677 | 1 | 3 |
| P27619 | 0 | 4 |
| Q94516 | 0 | 4 |
| Q9VI55 | 0 | 4 |
| A1ZBJ2 | 0 | 4 |
| P54367 | 0 | 4 |
| Q24246 | 0 | 4 |
| O97422 | 0 | 4 |
| Q7JQN4 | 0 | 4 |
| Q9VTB3 | 0 | 4 |
| M9PGI6 | 0 | 4 |
| Q9VWV5 | 0 | 4 |
| Q9V466 | 0 | 4 |
| Q9VPX5 | 0 | 4 |
| Q9V6B9 | 0 | 4 |
| Q9VV39 | 0 | 4 |
| Q9VFS5 | 0 | 4 |
| Q24276 | 0 | 4 |
| Q9VK06 | 0 | 4 |
| Q9VZF5 | 0 | 4 |
| P91929 | 0 | 4 |
| Q9VRH6 | 0 | 4 |
| Q9VXG0 | 0 | 4 |
| O46098 | 0 | 4 |
| Q9VKY3 | 0 | 4 |
| Q9V420 | 0 | 4 |
| M9MSL3 | 0 | 4 |
| Q9U3Z7 | 0 | 4 |
| Q7JV69 | 0 | 4 |

| | | |
|---|---|---|
| O17432 | 0 | 4 |
| Q9VLM8 | 2 | 1 |
| A0A0B4KER0 | 2 | 1 |
| Q9VEN9 | 2 | 1 |
| Q9VF87 | 1 | 2 |
| Q9VKK1 | 2 | 1 |
| O76876 | 1 | 2 |
| A1Z803 | 1 | 2 |
| Q9VTB0 | 1 | 2 |
| A0A0B4LHC9 | 2 | 1 |
| Q9VUL8 | 1 | 2 |
| Q9VM33 | 1 | 2 |
| Q9VBZ5 | 1 | 2 |
| Q9XZ53 | 1 | 2 |
| Q9V3E7 | 1 | 2 |
| Q9W335 | 1 | 2 |
| Q9VHR5 | 1 | 2 |
| Q8IMX4 | 1 | 2 |
| O97066 | 1 | 2 |
| Q9VYG1 | 1 | 2 |
| Q9W254 | 1 | 2 |
| Q9VRI0 | 1 | 2 |
| Q7K4H4 | 1 | 2 |
| Q8IMV6 | 1 | 2 |
| Q9VHW4 | 1 | 2 |
| Q9VC94 | 1 | 2 |
| E2QCS8 | 1 | 2 |
| Q9VSH0 | 1 | 2 |
| P18431 | 1 | 2 |
| A1Z7P5 | 1 | 2 |
| Q9VCQ7 | 2 | 1 |
| Q9VRE0 | 2 | 1 |
| E2QD16 | 2 | 1 |
| P45888 | 2 | 1 |
| Q9XZL8 | 2 | 1 |
| Q7KNA0 | 2 | 1 |
| A1Z9E2 | 2 | 1 |
| A1ZA22 | 2 | 1 |
| Q9W227 | 1 | 2 |
| Q7JX95 | 2 | 1 |
| P48612 | 1 | 2 |
| P49963 | 2 | 1 |
| Q9VIG0 | 1 | 2 |

| | | |
|---|---|---|
| Q9VSI1 | 1 | 2 |
| Q9VL69 | 1 | 2 |
| Q8SYG2 | 1 | 2 |
| Q7KNF2 | 1 | 2 |
| Q9Y171 | 1 | 2 |
| Q9VDS5 | 1 | 2 |
| Q9VEX9 | 1 | 2 |
| Q9I7U4 | 2 | 1 |
| Q9VX98 | 1 | 2 |
| Q9W5R5 | 2 | 1 |
| Q7JZF5 | 1 | 2 |
| Q9W124 | 2 | 1 |
| Q9VMV5 | 2 | 1 |
| Q9VXJ0 | 1 | 2 |
| Q7JZB4 | 2 | 1 |
| Q59DP9 | 2 | 1 |
| Q9V3P3 | 1 | 2 |
| Q9VCI7 | 1 | 2 |
| Q9VJZ4 | 1 | 2 |
| Q7JWI7 | 1 | 2 |
| P26019 | 2 | 0 |
| Q9VV61 | 1 | 2 |
| Q9VC06 | 1 | 2 |
| Q8SY39 | 1 | 2 |
| Q7KTL5 | 2 | 1 |
| Q8MSV2 | 1 | 2 |
| Q9W277 | 1 | 2 |
| Q9W547 | 1 | 2 |
| Q9VBP9 | 1 | 2 |
| Q7KTC0 | 1 | 2 |
| M9PEW1 | 0 | 3 |
| Q5U117 | 1 | 2 |
| Q9W0M6 | 1 | 2 |
| Q9VGU6 | 1 | 2 |
| Q9VAC1 | 1 | 2 |
| O61345 | 1 | 2 |
| Q9VXP3 | 2 | 1 |
| Q9VPT3 | 1 | 2 |
| Q9VLM5 | 1 | 2 |
| Q9VNX8 | 1 | 2 |
| Q9VJE5 | 2 | 1 |
| Q9U5L1 | 1 | 2 |
| Q7K231 | 1 | 2 |

| | | |
|---|---|---|
| Q9VK85 | 0 | 3 |
| Q9VZU8 | 0 | 3 |
| Q9VYV3 | 0 | 3 |
| Q9V6X7 | 0 | 3 |
| A1Z9G2 | 0 | 3 |
| P55162 | 2 | 1 |
| Q24050 | 0 | 3 |
| Q9VZE6 | 0 | 3 |
| Q9VPN5 | 0 | 3 |
| Q9VD00 | 0 | 3 |
| Q9V784 | 0 | 3 |
| Q7K126 | 0 | 3 |
| Q9VPQ2 | 0 | 3 |
| Q9VJ59 | 0 | 3 |
| Q9VC93 | 0 | 3 |
| P54351 | 0 | 3 |
| Q0E940 | 0 | 3 |
| Q9VNA3 | 0 | 3 |
| Q9W2S1 | 0 | 3 |
| Q7JVI6 | 0 | 3 |
| Q95RN0 | 0 | 3 |
| Q9VUN9 | 0 | 3 |
| Q9VBX3 | 0 | 3 |
| Q7JXF7 | 0 | 3 |
| Q9W4V8 | 0 | 3 |
| Q9VI57 | 0 | 3 |
| Q09332 | 0 | 3 |
| P92208 | 0 | 3 |
| Q9V3Y2 | 0 | 3 |
| Q9W1F7 | 0 | 3 |
| Q9VRP4 | 0 | 3 |
| Q7KVW1 | 0 | 3 |
| Q9VBS8 | 0 | 3 |
| Q8MT06 | 0 | 3 |
| F3YDF1 | 0 | 3 |
| Q9VXB0 | 0 | 3 |
| Q9VL10 | 0 | 3 |
| Q9VJ11 | 0 | 3 |
| Q9VJ43 | 0 | 3 |
| P16371 | 0 | 3 |
| Q9W3N1 | 0 | 3 |
| Q9VV89 | 0 | 3 |
| Q9V3P0 | 0 | 3 |

| | | |
|---|---|---|
| Q9GU50 | 0 | 3 |
| P49735 | 0 | 3 |
| Q7JWD3 | 0 | 3 |
| Q8IQV1 | 0 | 3 |
| Q9W552 | 0 | 3 |
| Q9VK63 | 0 | 3 |
| Q9VHR8 | 3 | 0 |
| A0A0B4KG96 | 0 | 3 |
| Q9VEN1 | 3 | 0 |
| Q9VZM5 | 0 | 3 |
| A0A4D6K5H4 | 0 | 3 |
| Q0KI05 | 2 | 0 |
| A1Z813 | 0 | 2 |
| A0A0B4K5Z8 | 0 | 2 |
| Q24318 | 0 | 2 |
| Q9W0R0 | 0 | 2 |
| M9PEI6 | 0 | 2 |
| Q9VIW3 | 0 | 2 |
| Q8IRH1 | 0 | 2 |
| Q9VC45 | 2 | 0 |
| Q9W2U7 | 0 | 2 |
| Q7KTG0 | 0 | 2 |
| Q9VUJ0 | 0 | 2 |
| P26686 | 0 | 2 |
| Q9VXE3 | 0 | 2 |
| Q9VFF3 | 0 | 2 |
| Q9VIX4 | 0 | 2 |
| A0A0B4K7C5 | 0 | 2 |
| A0A0B4KG70 | 0 | 2 |
| Q5U0V7 | 0 | 2 |
| Q94523 | 0 | 2 |
| Q9VWS3 | 2 | 0 |
| Q9VRJ2 | 2 | 0 |
| Q9VMQ7 | 0 | 2 |
| Q9VRJ7 | 0 | 2 |
| Q6NQY9 | 0 | 2 |
| Q9VRJ1 | 0 | 2 |
| P05990 | 0 | 2 |
| A0A0B4LH20 | 0 | 2 |
| Q9VS62 | 0 | 2 |
| Q9VJ87 | 0 | 2 |
| Q9XZ58 | 0 | 2 |
| P56079 | 0 | 2 |

| | | |
|---|---|---|
| Q8T0S6 | 0 | 2 |
| P51592 | 2 | 0 |
| Q7JUX9 | 0 | 2 |
| P39205 | 2 | 0 |
| Q9VSR5 | 0 | 2 |
| O46106 | 0 | 2 |
| Q7K511 | 0 | 2 |
| Q9VVM8 | 2 | 0 |
| P83094 | 2 | 0 |
| Q9W499 | 0 | 2 |
| Q9VMR6 | 0 | 2 |
| Q7K0F7 | 0 | 2 |
| A1ZAU8 | 0 | 2 |
| Q0E8J0 | 0 | 2 |
| Q9W0H8 | 0 | 2 |
| Q9V411 | 0 | 2 |
| Q7K3W2 | 0 | 2 |
| Q9GYU8 | 0 | 2 |
| Q9VBG6 | 0 | 2 |
| O02373 | 0 | 2 |
| A1Z992 | 0 | 2 |
| Q9VXQ0 | 0 | 2 |
| Q9W1H4 | 0 | 2 |
| Q7JRC9 | 0 | 2 |
| Q9VXK5 | 0 | 2 |
| Q9VZ82 | 0 | 2 |
| P35122 | 0 | 2 |
| Q9VJZ5 | 0 | 2 |
| Q9VPE2 | 2 | 0 |
| Q9VJ30 | 2 | 0 |
| Q9VBI2 | 0 | 2 |
| Q7K2E1 | 0 | 2 |
| P22465 | 0 | 2 |
| Q7JRE4 | 0 | 2 |
| Q9VMH2 | 0 | 2 |
| M9PBR6 | 0 | 2 |
| Q9VW19 | 0 | 2 |
| Q7KN61 | 0 | 2 |
| Q9VDE5 | 0 | 2 |
| Q9XZ34 | 2 | 0 |
| Q9W1G0 | 0 | 2 |
| O77477 | 0 | 2 |
| Q7JR49 | 0 | 2 |

| | | |
|---|---|---|
| A1Z6J5 | 0 | 2 |
| Q9W3N6 | 0 | 2 |
| Q9VC10 | 0 | 2 |
| Q9V428 | 0 | 2 |
| Q9VG80 | 0 | 2 |
| Q9VF28 | 0 | 2 |
| A1Z9A8 | 0 | 2 |
| Q9VZF6 | 0 | 2 |
| Q9VZZ9 | 0 | 2 |
| Q9W4A0 | 0 | 2 |
| Q9VIW7 | 0 | 2 |
| A1ZAN6 | 0 | 2 |
| Q9W3N9 | 0 | 2 |
| Q9VYT6 | 0 | 2 |
| P21914 | 0 | 2 |
| Q9VL25 | 0 | 2 |
| Q7PLS1 | 0 | 2 |
| Q7PLT4 | 0 | 2 |
| Q9VN88 | 0 | 2 |
| Q8SXX1 | 2 | 0 |
| Q24251 | 0 | 2 |
| Q9VQ89 | 0 | 2 |
| Q9VN39 | 0 | 2 |
| Q9VLQ1 | 0 | 2 |
| Q5BI62 | 2 | 0 |
| Q9VCF8 | 0 | 2 |
| Q7JWF1 | 0 | 2 |
| Q9V3F3 | 0 | 2 |
| P08646 | 0 | 2 |
| Q9VL02 | 0 | 2 |
| Q9V895 | 0 | 2 |
| Q9VF89 | 0 | 2 |
| Q6NN40 | 0 | 2 |
| A1ZA83 | 0 | 2 |
| Q7K1T1 | 0 | 2 |
| Q9VGZ3 | 0 | 2 |
| Q9VP29 | 0 | 2 |
| Q7K486 | 0 | 2 |
| Q9VNX4 | 0 | 2 |
| Q9VCB1 | 0 | 2 |
| X2JA38 | 0 | 2 |
| Q9VC87 | 0 | 2 |
| Q9VJ21 | 2 | 0 |

| | | |
|---|---|---|
| M9ND00 | 2 | 0 |
| Q9W0S2 | 0 | 2 |
| P35500 | 0 | 2 |
| Q9VGV8 | 0 | 2 |
| Q9VG81 | 0 | 2 |
| Q9VEJ0 | 0 | 2 |
| Q0E9H9 | 2 | 0 |
| Q9W366 | 2 | 0 |

**Table S2. dsRNA used in this study.**

| primer name | primer sequence |
|---|---|
| S1P-ds1F | TAATACGACTCACTATAGGGAAATCGGGAATCCATTTAGGG |
| S1P-ds1R | TAATACGACTCACTATAGGGTTTAGTTCGAGAGGAATGAC |
| S1P-ds2F | TAATACGACTCACTATAGGGGTAACGAACTGGAGAATTGTTC |
| S1P-ds2R | TAATACGACTCACTATAGGGATGTAAAGATCGGCGTCGGG |
| SREBP-ds1F | TAATACGACTCACTATAGGGGCGGTGTCAGTAGATTGTG |
| SREBP-ds1R | TAATACGACTCACTATAGGGAGTTCCTGGATATGGCTATTG |
| SREBP-ds2F | TAATACGACTCACTATAGGGAACGAGATGTCCGAGTGCAT |
| SREBP-ds2R | TAATACGACTCACTATAGGGGTTCAACCCAAGGTGAAGGA |
| ACLY-ds1F | TAATACGACTCACTATAGGGTGGGTACCTATCTCCTCCAA |
| ACLY-ds1R | TAATACGACTCACTATAGGGCTTCTGTGGTGGCAATGGT |
| ACLY-ds2F | TAATACGACTCACTATAGGGCTTCAGCGAAGCTCCACTCT |
| ACLY-ds2R | TAATACGACTCACTATAGGGAGCAAGCTGTTGAAGGAGGTC |
| ACS-ds1F | TAATACGACTCACTATAGGGCCCAGGCCGTTCCTAAC |
| ACS-ds1R | TAATACGACTCACTATAGGGCGAGGAGTACCAGAAGTTC |
| ACS-ds2F | TAATACGACTCACTATAGGGATCACTGAAGTGGCTCCATT |
| ACS-ds2R | TAATACGACTCACTATAGGGTGGCGGAGCGGATGTT |
| ACC-ds1F | TAATACGACTCACTATAGGGCTGCCTCTACGGCTTCC |
| ACC-ds1R | TAATACGACTCACTATAGGGCACCAGCCGGCTTATGA |
| ACC-ds2F | TAATACGACTCACTATAGGGCACACGCTCTCCTTCGTTTT |
| ACC-ds2R | TAATACGACTCACTATAGGGCTCTGAAGAAGCGGCAATTC |
| FASN1-ds1F | TAATACGACTCACTATAGGGAGTGCAAGTTACCGGGAATG |
| FASN1-ds1R | TAATACGACTCACTATAGGGCTCCAGTTCTCTGTACGCCTTG |
| FASN1-ds2F | TAATACGACTCACTATAGGGCCTGTCTTGGTGTAAGTGG |
| FASN1-ds2R | TAATACGACTCACTATAGGGAGCGTCGTCCTGGCAC |
| CG31523-ds1F | TAATACGACTCACTATAGGGTGGCGTTTGTTCGGTAAGTTTC |
| CG31523-ds1R | TAATACGACTCACTATAGGGATCGAATGGTGCCTACAAGG |
| CG31523-ds2F | TAATACGACTCACTATAGGGCCCCAGTATTGCAACCGTT |
| CG31523-ds2R | TAATACGACTCACTATAGGGCCATTTTCACCCACCAGTTC |
| CG31522-ds1F | TAATACGACTCACTATAGGGCGTCTTCCGCTGTTTCAG |
| CG31522-ds1R | TAATACGACTCACTATAGGGCCCTGGCCAACGGACAC |
| CG31522-ds2F | TAATACGACTCACTATAGGGTTACTGTTTGTCGCCGACTG |
| CG31522-ds2R | TAATACGACTCACTATAGGGCACCGCATCAAATGCACTAA |
| CG3971-ds1F | TAATACGACTCACTATAGGGCATTCAGGGGCTAGTGGAAA |
| CG3971-ds1R | TAATACGACTCACTATAGGGCACCAGCACCTAACTCCCAT |
| CG3971-ds2F | TAATACGACTCACTATAGGGAAGCGGGCCGCCTTC |
| CG3971-ds2R | TAATACGACTCACTATAGGGTCACCACATCACCGTGCT |
| Desat1-ds1F | TAATACGACTCACTATAGGGCTAAGATGCACGTCTGCCAC |
| Desat1-ds1R | TAATACGACTCACTATAGGGCCCAGTCCATCTCAGACTCAC |
| Desat1-ds2F | TAATACGACTCACTATAGGGAAGCAATCGGCATCAAAATC |
| Desat1-ds2R | TAATACGACTCACTATAGGGCACGCGTGGTTTCTTCTTCT |
| CG9743-ds1F | TAATACGACTCACTATAGGGCGGCTTGTTGCCGTACA |

| | |
|---|---|
| CG9743-ds1R | TAATACGACTCACTATAGGGTCACCGTTTGGGTCTCCG |
| CG9743-ds2F | TAATACGACTCACTATAGGGTTGGGTGATGGGTCATAGCC |
| CG9743-ds2R | TAATACGACTCACTATAGGGATCCCCTGACATGGTCTTGA |
| Gpat4-ds1F | TAATACGACTCACTATAGGGCAGCCGATTGGCAAACTC |
| Gpat4-ds1R | TAATACGACTCACTATAGGGCGGCGACGCCTTCTG |
| mino-ds1F | TAATACGACTCACTATAGGGCAGATGGCTTGTAGACCTTG |
| mino-ds1R | TAATACGACTCACTATAGGGAACCTGCAAATTCCGGTCT |
| Agpat2-ds1F | TAATACGACTCACTATAGGGCTCGGGTAGGATGTGGATTA |
| Agpat2-ds1R | TAATACGACTCACTATAGGGTCCATCATGGTGCTTCCG |
| Agpat3-ds1F | TAATACGACTCACTATAGGGAGTCCCAGGACGGAAAGT |
| Agpat3-ds1R | TAATACGACTCACTATAGGGAGTTTGCTGCATGGAAAGAG |
| Mdy-ds1F | TAATACGACTCACTATAGGGAAGCCCAAACGCAGACC |
| Mdy-ds1R | TAATACGACTCACTATAGGGAACCGCAAGTCAACACAAAA |
| app-ds1f | TAATACGACTCACTATAGGGAGTGGTGAATATTTCAAGCAG |
| app-ds1r | TAATACGACTCACTATAGGGTGGTCCTGCTGGTGGTTC |
| CG1407-ds1f | TAATACGACTCACTATAGGGTCGCAAGCTGTTAACAAACG |
| CG1407-ds1r | TAATACGACTCACTATAGGGAACCAATTTGAGTTTCCCCA |
| CG5196-ds1f | TAATACGACTCACTATAGGGAATGTATACTGATCACAACCCT |
| CG5196-ds1r | TAATACGACTCACTATAGGGTCGCTGCGTCAAGAAGA |
| CG5880-ds1f | TAATACGACTCACTATAGGGCTCGAAGGGAGCATCATTTA |
| CG5880-ds1r | TAATACGACTCACTATAGGGCCGAATGAGTATGATGAATTTG |
| CG8314-ds1f | TAATACGACTCACTATAGGGAGTACACATCTTCACCCGGC |
| CG8314-ds1r | TAATACGACTCACTATAGGGAAGAGCATCCAGTCGGTGTT |
| CG10344-ds1f | TAATACGACTCACTATAGGGCAGGTAGGAACTGCACAGCA |
| CG10344-ds1r | TAATACGACTCACTATAGGGCTGCTGCATTGGACTCAGAA |
| CPT2-ds1f | TAATACGACTCACTATAGGGCGCCAGCAAATGCTTGA |
| CPT2-ds1r | TAATACGACTCACTATAGGGCAGGTGGATGAAGGTGGC |
| Dnz1-ds1f | TAATACGACTCACTATAGGGTGGTGGTCAGAATGATCCAA |
| Dnz1-ds1r | TAATACGACTCACTATAGGGTGAAACGCTCGATTTGTGAA |
| GABPI-ds1f | TAATACGACTCACTATAGGGATGATCGCGCCTCTTCA |
| GABPI-ds1r | TAATACGACTCACTATAGGGCTGGTGGGCTTCACCATC |
| Hip14-ds1f | TAATACGACTCACTATAGGGCCCTTCCACGACTTAAGGA |
| Hip14-ds1r | TAATACGACTCACTATAGGGTCAAGTCAAAGGCCTCGTT |
| lace-ds1f | TAATACGACTCACTATAGGGTATTTCTTTTTCAGAGCAATAACC |
| lace-ds1r | TAATACGACTCACTATAGGGACACGTTGCTTAAATCTGGG |
| por-ds1f | TAATACGACTCACTATAGGGTTCGTTTGTTCCAGTTTCCC |
| por-ds1f | TAATACGACTCACTATAGGGATGTCGGTATAGCCACTGCC |
| rasp-ds1f | TAATACGACTCACTATAGGGCACATGTCCGAGTAGAAGTG |
| rasp-ds1r | TAATACGACTCACTATAGGGACAGATCGCGTTGGTTACAC |
| Spt-I-ds1f | TAATACGACTCACTATAGGGCATTAAAGTGCTCCGTAACTC |
| Spt-I-ds1r | TAATACGACTCACTATAGGGCGCAAGTACGGAGTTGGA |
| whd-ds1f | TAATACGACTCACTATAGGGTTCCTGGTGCTCGATAAGT |
| whd-ds1r | TAATACGACTCACTATAGGGCAAGCTGCTGCTCATGTAC |

| | |
|---|---|
| Ykt6-ds1f | TAATACGACTCACTATAGGGCTCTGCAGCGACAATTTC |
| Ykt6-ds1r | TAATACGACTCACTATAGGGCGAGGCGCGTCTCCTG |
| Zdhhc8-ds1f | TAATACGACTCACTATAGGGCTAAAGAAGGTTGAAAATTTGC |
| Zdhhc8-ds1r | TAATACGACTCACTATAGGGCCACCCGCCAGCCAAG |
| Nmt-ds1f | TAATACGACTCACTATAGGGCATGTCCTTCGCTGTGATT |
| Nmt-ds1r | TAATACGACTCACTATAGGGAACTATGTGGAGGACGATGA |
| Nmt-ds2f | TAATACGACTCACTATAGGGAAATATCGAAGCTTGGCCGT |
| Nmt-ds2r | TAATACGACTCACTATAGGGTAAGCTCACATTCTCGCCCT |
| Arf1-ds1f | TAATACGACTCACTATAGGGTAGCGATTAGCGTTCTTCA |
| Arf1-ds1r | TAATACGACTCACTATAGGGCTGCCAAATGCAATGAACG |
| Arf1-ds2f | TAATACGACTCACTATAGGGCACATTGAAACCTAGGGG |
| Arf1-ds2r | TAATACGACTCACTATAGGGACTACCGAGCATCCGAAAGA |
| alphaCOP-ds1f | TAATACGACTCACTATAGGGAGGAAGCTAAGCTTGTCAAA |
| alphaCOP-ds1r | TAATACGACTCACTATAGGGACGAGTCTGGAGTGTTTATC |
| betapCOP-ds1f | TAATACGACTCACTATAGGGCTCTTTAACCAGAGACATGTT |
| betapCOP-ds1r | TAATACGACTCACTATAGGGTTTCGGAGTGCGTCAAAAC |
| betaCOP-ds1f | TAATACGACTCACTATAGGGACTCTGGGTGGCATAGGTT |
| betaCOP-ds1f | TAATACGACTCACTATAGGGACACTGGCAAGTACAGGCAG |
| deltaCOP-ds1f | TAATACGACTCACTATAGGGCCGCCTTGGATTTGGTGT |
| deltaCOP-ds1r | TAATACGACTCACTATAGGGTCCCGAGTACAGCCACT |
| epsilonCOP-ds1f | TAATACGACTCACTATAGGGAGGTGCCAGATGTTGGTCTC |
| epsilonCOP-ds1r | TAATACGACTCACTATAGGGCCAACTCGGTGCTATTCGAT |
| gammaCOP-ds1f | TAATACGACTCACTATAGGGTCCATCTGGCAACGACCC |
| gammaCOP-ds1r | TAATACGACTCACTATAGGGTCGGTAGAGCGTCTGATG |
| zetaCOP-ds1f | TAATACGACTCACTATAGGGCCGTCGCAGATCTCGTC |
| zetaCOP-ds1r | TAATACGACTCACTATAGGGCATCCTGGCCAAGTACTAC |
| Chc-ds1f | TAATACGACTCACTATAGGGCACAATCCACGCTCGTAG |
| Chc-ds1r | TAATACGACTCACTATAGGGCCCGAACGGGTGAAGAAC |
| Clc-ds1f | TAATACGACTCACTATAGGGTTTCCCGCCTTGTTCACC |
| Clc-ds1r | TAATACGACTCACTATAGGGAGTCAGCTCTCGGGGAC |
| AP-1-2beta-ds1f | TAATACGACTCACTATAGGGAATCCTTCAAGGAAGCTGTC |
| AP-1-2beta-ds1r | TAATACGACTCACTATAGGGGCCCTGCGCAACATCAA |
| Gga-ds1f | TAATACGACTCACTATAGGGTGGAACAGCTCCTCGCT |
| Gga-ds1r | TAATACGACTCACTATAGGGCCAATTGCGCTCTACTGG |
| Sar1-ds1f | TAATACGACTCACTATAGGGTTGTTAGCTGATACAGTCCG |
| Sar1-ds1r | TAATACGACTCACTATAGGGACGCGTCTGGAAGGAC |
| Sec13-ds1f | TAATACGACTCACTATAGGGAATGTGTGCAGCACAGTCG |
| Sec13-ds1r | TAATACGACTCACTATAGGGACAGCAACCACGACTCCT |
| Sec23-ds1f | TAATACGACTCACTATAGGGAATGGGGGCTGCATGC |
| Sec23-ds1r | TAATACGACTCACTATAGGGACGACGAGCTGAAGCAC |
| Sec24AB-ds1f | TAATACGACTCACTATAGGGTCACGGAAGGGATGTATCAC |
| Sec24AB-ds1r | TAATACGACTCACTATAGGGAGCCCAGCCCATGATGC |
| Sec24CD-ds1f | TAATACGACTCACTATAGGGCCTTATCAGTTCCCAGAAGC |

| | |
|---|---|
| Sec24CD-ds1r | TAATACGACTCACTATAGGGTTCCCGCCTTTATCTTCATC |
| Sec31-ds1f | TAATACGACTCACTATAGGGCATGGTAAACCACTTGCGTG |
| Sec31-ds1r | TAATACGACTCACTATAGGGTAATGGTTTCGTGTGGCAAA |
| Sccpdh2-ds1f | TAATACGACTCACTATAGGGAAGGAGAACTTGGCCATAATG |
| Sccpdh2-ds1r | TAATACGACTCACTATAGGGGAGCGTGGCGTGTACG |
| Sccpdh2-ds2f | TAATACGACTCACTATAGGGTGGCCGCCAGCTGCTGCT |
| Sccpdh2-ds2r | TAATACGACTCACTATAGGGCTATTTGTTGGCCACGATTTCAAAC |
| CG33303-ds1f | TAATACGACTCACTATAGGGTGTCACTGGATCCCTCAGG |
| CG33303-ds1r | TAATACGACTCACTATAGGGCCAGGAGCCACTAGTCATC |
| CG5854-ds1f | TAATACGACTCACTATAGGGTTATCGCCAATGCCATAGAC |
| CG5854-ds1r | TAATACGACTCACTATAGGGATGCAAGGCGGCTTTTG |
| Droj2-ds1f | TAATACGACTCACTATAGGGCATCGAGCGTTTTAACGATT |
| Droj2-ds1r | TAATACGACTCACTATAGGGTGGAGGAGCTGTACAATGG |
| Gfat2-ds1f | TAATACGACTCACTATAGGGCACCCAGAACCACGGTCT |
| Gfat2-ds1r | TAATACGACTCACTATAGGGTCCAGGCGAGGCTCAC |
| Gp93-ds1f | TAATACGACTCACTATAGGGTCCTTGTCGATCTTCTT |
| Gp93-ds1r | TAATACGACTCACTATAGGGAGGAGGCCGAAGATGAGA |
| Hel25E-ds1f | TAATACGACTCACTATAGGGTATGTAGAGAGATCGATTTCCT |
| Hel25E-ds1r | TAATACGACTCACTATAGGGTCATCTTTGTGAAGTCTGTGC |
| Mcad-ds1f | TAATACGACTCACTATAGGGCAACGCTGGGCCAAACC |
| Mcad-ds1r | TAATACGACTCACTATAGGGTGATCCTGTCCGGTAACAAG |
| Nfs1-ds1f | TAATACGACTCACTATAGGGAAGAAAGTTCCGCAGCAG |
| Nfs1-ds1r | TAATACGACTCACTATAGGGCGTTTCTACGGCACTAAGA |
| Rab5-ds1f | TAATACGACTCACTATAGGGTCCTGGCCAGCCGTGT |
| Rab5-ds1r | TAATACGACTCACTATAGGGCAACCACTCCACGCAGC |
| Raf-ds1f | TAATACGACTCACTATAGGGAGGTCTGGCCTGCATTATC |
| Raf-ds1r | TAATACGACTCACTATAGGGCATGTGGAGGAGATCTTTGTC |
| RpS10b-ds1f | TAATACGACTCACTATAGGGTGCCGCCGGGAGCAC |
| RpS10b-ds1r | TAATACGACTCACTATAGGGCGCTGAAGCGCCCTG |
| cer-ds1f | TAATACGACTCACTATAGGGAATTTGGCGGCACCTTTTT |
| cer-ds1r | TAATACGACTCACTATAGGGAAGTTCGACAAGAACTACGA |
| mEFTu1-ds1f | TAATACGACTCACTATAGGGTGAAGCTATCCACCTCCTGG |
| mEFTu1-ds1r | TAATACGACTCACTATAGGGCACTACGGCCACACAGATTG |
| sav-ds1f | TAATACGACTCACTATAGGGCACATGCGGATGGGAGG |
| sav-ds1r | TAATACGACTCACTATAGGGAACTATCTAACGATCCTGCTG |
| wts-ds1f | TAATACGACTCACTATAGGGCAGCCCGTTCTGGGTTATTA |
| wts-ds1r | TAATACGACTCACTATAGGGACGTGTGCCAAAGATGATGAC |
| pcb-ds1f | TAATACGACTCACTATAGGGCTTTCCAGCGGCTTCCA |
| pcb-ds1r | TAATACGACTCACTATAGGGAACCACCGAAGGGACTCC |
| Pdi-ds1f | TAATACGACTCACTATAGGGAGCGATAAGTAAGCGGCAAA |
| Pdi-ds1r | TAATACGACTCACTATAGGGACTTCTTAAGTTTCGGCGCA |
| PH4alphaEFB-ds1f | TAATACGACTCACTATAGGGGAGAAAGTTCAGCTTGTCCT |
| PH4alphaEFB-ds1r | TAATACGACTCACTATAGGGGACTGGCGATTGATGCTC |

| | |
|---|---|
| Tom70-ds1f | TAATACGACTCACTATAGGGTAGCTATAATAGCATCAAAGTCC |
| Tom70-ds1r | TAATACGACTCACTATAGGGGCCTCGCTGGAGTTCAA |
| Cyp6a20-ds1f | TAATACGACTCACTATAGGGCAATGACCTTTTCCAAATAAGTC |
| Cyp6a20-ds1r | TAATACGACTCACTATAGGGCTTCGGCTTGGATTGTCATA |
| EMC1-ds2f | TAATACGACTCACTATAGGGAAACGTAGAAGGTGCCGCTA |
| EMC1-ds2r | TAATACGACTCACTATAGGGCTTGAGTGGGAGTGGTCCAT |
| Sec63-ds1f | TAATACGACTCACTATAGGGGTCACCACATTGGTGTTCTC |
| Sec63-ds1r | TAATACGACTCACTATAGGGCTGTGCCGCATGTACTCC |
| CaBP1-ds1f | TAATACGACTCACTATAGGGGAATGTGTGGCAGAACGGA |
| CaBP1-ds1r | TAATACGACTCACTATAGGGGAAGAAGGTGCAGGGTGTC |
| P58IPK-ds1f | TAATACGACTCACTATAGGGCAGGCAGCCAATTTCTTCTC |
| P58IPK-ds1r | TAATACGACTCACTATAGGGCCCATCCCTAGAGAACGTGA |
| Hrd3-ds1f | TAATACGACTCACTATAGGGTCATTATCGGCCCTGATTTG |
| Hrd3-ds1r | TAATACGACTCACTATAGGGGATTCCTTTATTCGGTGGGC |
| Oatp58Db-ds1f | TAATACGACTCACTATAGGGCAGAGAATCCCAACGCCC |
| Oatp58Db-ds1r | TAATACGACTCACTATAGGGGGATCTTTATTTTATTTTCACTTCC |
| Ufl1-ds1f | TAATACGACTCACTATAGGGAATGCCATCAAGATAAACAAAG |
| Ufl1-ds1r | TAATACGACTCACTATAGGGCCTGGCCGCCATTACCA |
| GlcAT-I-ds1f | TAATACGACTCACTATAGGGCCTGCGGATTCCTGATGA |
| GlcAT-I-ds1r | TAATACGACTCACTATAGGGGCTGCTGCCGCACCT |
| Sec22-ds1f | TAATACGACTCACTATAGGGTCCTTCCACTGGCTGCTACT |
| Sec22-ds1r | TAATACGACTCACTATAGGGACCGGTGACGTGGTTACATT |
| CG6263-ds1f | TAATACGACTCACTATAGGGTCTTCGCTGGGGGTCTT |
| CG6263-ds1r | TAATACGACTCACTATAGGGACGGGCACACTCACACAA |
| Cdc50-ds2f | TAATACGACTCACTATAGGGATTCGGCCTTCAAGCAACAG |
| Cdc50-ds2r | TAATACGACTCACTATAGGGATTGAAGTCCGAGGGCAGG |
| Sc2-ds2f | TAATACGACTCACTATAGGGACCCTGTTAAACGCTAGCCA |
| Sc2-ds2r | TAATACGACTCACTATAGGGAAACTTTTCGGTGCACATTGC |
| CG8507-ds1f | TAATACGACTCACTATAGGGTATTTCTCCTTTTGCAAAGCAT |
| CG8507-ds1r | TAATACGACTCACTATAGGGAAGACTTGGACACCTACAATCTG |
| NTPase-ds1f | TAATACGACTCACTATAGGGACTCGATGCACGTCTAAGGC |
| NTPase-ds1r | TAATACGACTCACTATAGGGAGATCTGTGCCATTCCCAAC |
| Vap33-ds1f | TAATACGACTCACTATAGGGCCTCGCTGGAGAGCTTC |
| Vap33-ds1r | TAATACGACTCACTATAGGGCCGAGCAGCTGATGGAC |
| TER94-ds2f | TAATACGACTCACTATAGGGCTACCGACAATGCACGAATG |
| TER94-ds2r | TAATACGACTCACTATAGGGTACACCGCAAAACAAAACCA |
| ERp60-ds1f | TAATACGACTCACTATAGGGCCAGCTCCTCGTAGATGG |
| ERp60-ds1r | TAATACGACTCACTATAGGGGTGGGACACCGCACTCA |
| Past1-ds1f | TAATACGACTCACTATAGGGCGTTACGCGGCAGGG |
| Past1-ds1r | TAATACGACTCACTATAGGGGTACGGCAACGCCTTCC |
| CG6766-ds1f | TAATACGACTCACTATAGGGAAGGGGCGTCAGATCAGT |
| CG6766-ds1r | TAATACGACTCACTATAGGGATACAGCGATGGCACTATGT |
| Mitofilin-ds1f | TAATACGACTCACTATAGGGCGCAGCAGCTTATCAGAC |

| | |
|---|---|
| Mitofilin-ds1r | TAATACGACTCACTATAGGGCGAGATCGCCCTGAGTG |
| Gp93-ds2f | TAATACGACTCACTATAGGGCGAAGGTGCTCTTTAGCGAC |
| Gp93-ds2r | TAATACGACTCACTATAGGGTACTACATCGCTGGTGCCAA |
| bai-ds1f | TAATACGACTCACTATAGGGCGATGATCTGGCCGGGT |
| bai-ds1r | TAATACGACTCACTATAGGGATGGCCTGTGCCTGGAG |
| Hmu-ds1f | TAATACGACTCACTATAGGGAACTCCAGGACGTGTGATACC |
| Hmu-ds1r | TAATACGACTCACTATAGGGGCGATGCGTCTGACAAAGTA |
| Ant2-ds1f | TAATACGACTCACTATAGGGTACTTCTTCATCTCGTCGTAA |
| Ant2-ds1r | TAATACGACTCACTATAGGGGCGGCAATCGGGAGTTC |
| CG1309-ds1f | TAATACGACTCACTATAGGGGTAGTTTCGAATGGCTACCTT |
| CG1309-ds1r | TAATACGACTCACTATAGGGGCTGATCTCCGGACGC |
| Mec2-ds1f | TAATACGACTCACTATAGGGCTCGGCGGCGATCAC |
| Mec2-ds1r | TAATACGACTCACTATAGGGGAGCGGGCGATCATCTT |
| Clic-ds1f | TAATACGACTCACTATAGGGTGCAGGCGCGGCATC |
| Clic-ds1r | TAATACGACTCACTATAGGGCCACACACCCGCCCATC |
| ATP6AP2-ds1f | TAATACGACTCACTATAGGGCTTCTTGATGCGGGTAGAGG |
| ATP6AP2-ds1r | TAATACGACTCACTATAGGGAAGCACCTGAACCCCTCTCT |
| Jwa-ds1f | TAATACGACTCACTATAGGGATGGGCGTGGAGGGAG |
| Jwa-ds1r | TAATACGACTCACTATAGGGCGCCTCTGCGAACCCT |
| Tango5-ds1f | TAATACGACTCACTATAGGGGCGCGACAGCAAATTCC |
| Tango5-ds1r | TAATACGACTCACTATAGGGCCAATCCCCTATTCGACCT |
| ttm50-ds1f | TAATACGACTCACTATAGGGGTCGCGATTAAGGTTGTCC |
| ttm50-ds1r | TAATACGACTCACTATAGGGGTGGCTGCTGGATTCTGG |
| Dpck-ds1f | TAATACGACTCACTATAGGGAGGAAGCTGATGCGGTTC |
| Dpck-ds1r | TAATACGACTCACTATAGGGGTGCTGGCGGCAGATTC |
| CG10470-ds1f | TAATACGACTCACTATAGGGTTTTGGATGTTCTGACGAATG |
| CG10470-ds1r | TAATACGACTCACTATAGGGGTGATGGTTTACCGGACGG |
| spidey-ds1f | TAATACGACTCACTATAGGGTGGAACTGTACACGCTCAG |
| spidey-ds1r | TAATACGACTCACTATAGGGCGGTGTGGAGGTGCGT |
| PIG-B-ds1f | TAATACGACTCACTATAGGGCTCTGTATTCCCGAGCTATT |
| PIG-B-ds1r | TAATACGACTCACTATAGGGCGTTGCTGGTCTGGGAATA |
| Cds-ds1f | TAATACGACTCACTATAGGGATTCCTGGGGCGTGAACA |
| Cds-ds1r | TAATACGACTCACTATAGGGCTGAAGTTCCTGGTGACCTA |
| MAN1-ds1f | TAATACGACTCACTATAGGGATCCAGCGACTATTAAGAGAA |
| MAN1-ds1r | TAATACGACTCACTATAGGGGTTCTTAATATTTTTGACAATAATCAC |
| CG33523-ds1f | TAATACGACTCACTATAGGGATGACTCCCAAAGATGATAGTTA |
| CG33523-ds1r | TAATACGACTCACTATAGGGTTTCCATCCGACCGATATTG |
| VhaAC45-ds1f | TAATACGACTCACTATAGGGGAGTCGCCGAAGGGAAAG |
| VhaAC45-ds1r | TAATACGACTCACTATAGGGCGCCATAGCTGCTATCAG |
| PIG-S-ds1f | TAATACGACTCACTATAGGGCAACAGTTTGATGAGGCTC |
| PIG-S-ds1r | TAATACGACTCACTATAGGGGCGCGACCAGAGCAAA |
| CG5196-ds2f | TAATACGACTCACTATAGGGCAGAGTAAACGGGATGACGG |
| CG5196-ds2r | TAATACGACTCACTATAGGGAGCTAGCTGGCGTCAAGAAG |

| | |
|---|---|
| p24-1-ds1f | TAATACGACTCACTATAGGGCACCATAATCTGGACGAGAC |
| p24-1-ds1r | TAATACGACTCACTATAGGGCTTCGATCTGGCCGATAAC |
| wol-ds1f | TAATACGACTCACTATAGGGGAAACTCCTTGTTGCAATCG |
| wol-ds1r | TAATACGACTCACTATAGGGCTTCTATGCTGGACGAGTG |
| EMC7-ds1f | TAATACGACTCACTATAGGGCATCCGGGTGGTGGACAT |
| EMC7-ds1r | TAATACGACTCACTATAGGGGCCTAAAGTTATTTGTTTTTACG |
| tweek-ds1f | TAATACGACTCACTATAGGGTGTGATGCTCCGAGAGTACG |
| tweek-ds1r | TAATACGACTCACTATAGGGGCAACTCGGGAGCTTTAGTG |
| para-ds1f | TAATACGACTCACTATAGGGCGGGCCGCTATCTCAC |
| para-ds1r | TAATACGACTCACTATAGGGGTTGGGATGGTGTACTGGAC |
| Afg3l2-ds1f | TAATACGACTCACTATAGGGGACGTTTTCAGGTTACCCAG |
| Afg3l2-ds1r | TAATACGACTCACTATAGGGCCGCAGCAGTATATTGATTTA |
| Tango1-ds1f | TAATACGACTCACTATAGGGGCCTTCCTTTGAGAAGGTG |
| Tango1-ds1r | TAATACGACTCACTATAGGGTGCAGCGCTCAAGTGAG |
| Tango1-ds2f | TAATACGACTCACTATAGGGCAGAGGCGTTAAGAACAGGC |
| Tango1-ds2r | TAATACGACTCACTATAGGGATTGCCAAAATCACCTACGC |
